# AI-generated small binder improves prime editing

**DOI:** 10.1101/2024.09.11.612443

**Authors:** Ju-Chan Park, Heesoo Uhm, Yong-Woo Kim, Ye Eun Oh, Sangsu Bae

## Abstract

The prime editing 2 (PE2) system comprises a nickase Cas9 fused to a reverse transcriptase utilizing a prime editing guide RNA (pegRNA) to introduce desired mutations at target genomic sites. However, the PE efficiency is limited by mismatch repair (MMR) that excises the DNA strand containing desired edits. Thus, inhibiting key components of MMR complex through transient expression of a dominant negative MLH1 (MLH1dn) exhibited approximately 7.7-fold increase in PE efficiency over PE2, generating PE4. Herein, by utilizing a generative artificial intelligence (AI) technologies, RFdiffusion and AlphaFold 3, we ultimately generated a de novo MLH1 small binder (named MLH1-SB), which bind to the dimeric interface of MLH1 and PMS2 to disrupt the formation of key MMR components. MLH1-SB’s small size (82 amino acids) allowed it to be integrated into pre-existing PE architectures via the 2A system, creating a novel PE-SB platform. Resultantly, by incorporating MLH1-SB into PE7, we have developed an improved PE architecture called PE7-SB, which demonstrates the highest PE efficiency to date (29.4-fold over PE2 and 2.4-fold over PE7 in HeLa cells), providing an insight that generative AI technologies will boost up the improvement of genome editing tools.

## Introduction

Since the advent of CRISPR-Cas9 genome editing technology ^1–4^, efforts have been made to repurpose this system, which has led to the development of base editors (BEs) and prime editors ^5–7^. While BEs are capable of converting one or a few nucleotides, prime editors enable the introduction of base substitutions and small insertion and deletion (indel) mutations ^8,9^. Due to the flexibility and precision, prime editors are attracting considerable attention for therapeutic applications such as cell and gene therapy as well as disease modeling ^10,11^. Therefore, various attempts have been made to enhance the efficiency of prime editing (PE), thus far.

The initial optimized PE architecture, PE2, consists of nickase Cas9 (nCas9) fused to reverse transcriptase (RT), synthesizing a DNA strand that contains the desired mutation at the target site ^7^. This process utilizes the 3′ extension of the prime editing guide RNA (pegRNA) as a template for reverse transcription. To increase PE efficiency, PE3 or PE5 employs an additional nicking guide RNA (ngRNA) to generate a second nick at the non-edited strand, altering the flap equilibrium to include a more desired edit ^7,12^. Further research revealed that the incorporation of the desired edits at the target site is impeded by mismatch repair (MMR) pathway ^12^. In MMR, MutS homologs (MSH2–MSH6 complex for MutSα and MSH2–MSH3 complex for MutSβ) recognize the mismatch and recruit the MutLα complex (composed of MLH1 and PMS2) to promote the progression of MMR ^13^. Thus, inhibiting these key components of MMR (e.g., MSH2, MSH3, MSH6, MLH1, and PMS2) substantially enhances PE efficiency ^12,14,15^. Notably, the co-delivery of a dominant negative MLH1 protein (MLH1dn) to inhibit the MutLα complex, along with prime editors, is defined as the PE4 architecture ^12^. RNA pseudoknot motifs are also incorporated to the 3′ terminus of pegRNAs to enhance their stability and prevent degradation of the 3′ extension ^16^. Moreover, additional protein evolution using a phage-assisted continuous evolution further enhances PE efficiency, generating various PE6 architectures ^17^. The PE efficiency can be further enhanced by protecting the 3′ end of pegRNA with RNA-binding exonuclease protection factor La, defined as the PE7 architecture ^18^. Despite the continuous efforts to upgrade PE activity, it is still unsatisfactory especially for therapeutic applications, particularly in the absence of ngRNAs that can generate undesired indel mutations.

Recent advances in artificial intelligence (AI) technologies have enabled the prediction of sequence-based protein structures ^19–22^ as well as the generation of de novo proteins/peptides ^23^. Moreover, these technologies have been applied to the development of the CRISPR toolbox ^24–26^. However, the utilization of AI technologies in advancing CRISPR genome editing tools has been primarily focused on enhancing the catalytic activity of key enzymes, such as Cas9 ^24^ and TadA ^26^. In addition, the application of RFdiffusion ^23^, an AI technology for binder protein generation, is primarily restricted to the development of single-site binders ^23^ and replacing antibodies ^27^. In this study, we harnessed RFdiffusion to inhibit the MMR pathway to enhance PE efficiency. Importantly, the success rate was exceptionally high through the use of two new approaches: i) using multiple binding sites to design de novo proteins including a non-competitive anchoring site and an inhibiting site against critical activity beside a bare competitive binding site, and ii) developing a new AlphaFold 3 ^20^-based filtering system to choose high-score candidates. Through this, the RFdiffusion-generated MLH1 small binders (named MLH1-SBs) significantly improved the PE efficiency and were compatible with pre-existing PE architectures including PE2, PEmax and PE7 ^7,12,18^. We ultimately selected a MLH1-SB with the size of 82 amino acids. The MLH1-SB’s small size allowed it to be integrated into PE2 and PE7 architectures, generating PE2-SB and PE7-SB, respectively. PE7-SB exhibited the highest PE efficiency, approximately 29- and 2.4-fold increase compared to PE2 and PE7, respectively.

## Results

### Generating de novo proteins binding to multiple hotspots of MLH1 using RFdiffusion

The incorporation of a PE-synthesized DNA strand that contains desired edits into the target genome can be interrupted by the MMR pathway. In this process, mismatch-recognizing MutS homologs recruit MutLα complex ^28,29^ for further progression, ultimately lowering the PE efficiency (**Fig. 1A**). To inhibit this process, we decided to generate de novo small binders by RFdiffusion, which bind to the dimeric interface of MLH1 and PMS2 to disrupt the formation of the MutLα complex (i.e., MLH1 C-terminal and PMS2 C-terminal) (**Fig. 1B**). However, although each structure of MLH1 and PMS2 was deposited in the Protein Data Bank (PDB), the complex structure of two has not been revealed yet. Thus, we modeled the dimeric interface structure using AlphaFold 3 to obtain the complex formation between MLH1 C-terminal and PMS2 C-terminal (**Fig. 1C**). Until now, most previous binder design strategies have focused on a single binding target site, although multiple hotspots were considered for binder design ^23,30–33^. But, we assumed that multiple binding sites from a single binder could simultaneously inhibit multiple activities and thereby increase the overall inhibition efficacy. Thus, we used three different binding sites in MLH1 to increase binder’s MMR inhibition efficacy. First, W538 and Q542 correspond to the “Main binding” site between MLH1 and PMS2 in which a binder would hinder the complex formation. Y750, located in the nuclease activity domain of the MutLα complex, corresponds to the “Critical activity” site ^28^. M682 is a hotspot for “Additional binding” site that provides a non-competitive anchoring site for binders, and thus further facilitates the inhibition activity of the binder designs (**Fig. 1C**). As a result, total 1978 binder candidates were generated using the “Main binding” and “Critical activity” hotspots (named “MC” binders), (**Figs. 1D** and **S1A**), and total 821 candidates were generated using the “Main binding,” “Critical activity,” and “Additional binding” hotspots (named “MCA” binders) (**Figs. 1D** and **S1B**).

**Figure 1.**
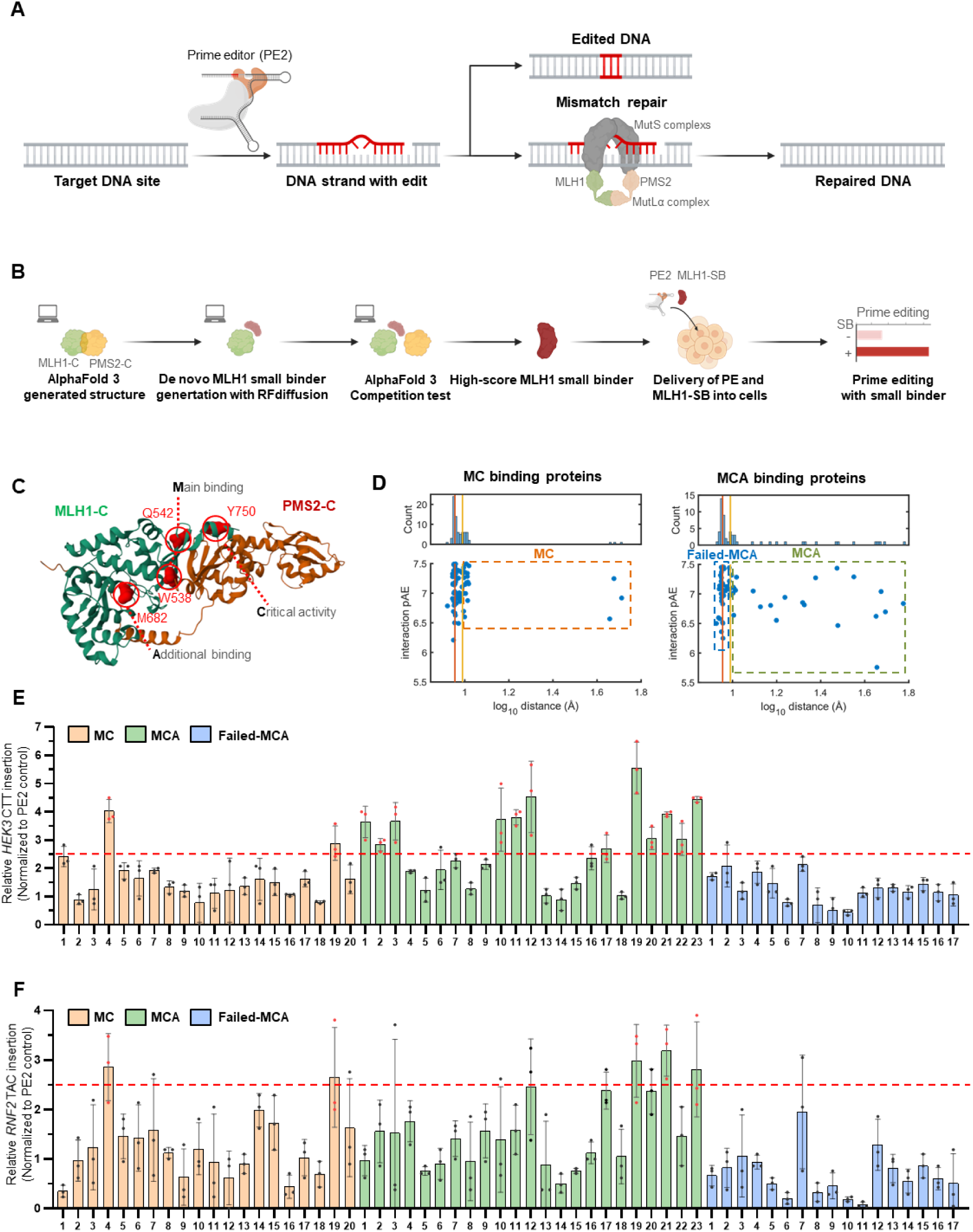
AI-generated MLH1 small binders improves prime editing. (A) Scheme of prime editing. DNA and RNA with intended mutation is colored in red. Created with BioRender.com (B) Scheme of MLH1 small binder generation. MLH1-C, C-terminal domain of MLH1; PMS2-C; C-terminal domain of PMS2, MLH1-SB; MLH1 small binder. Created with BioRender.com (C) Predicted structure of dimeric interface of MLH1 and PMS2 by AlphaFold 3. Main binding (W538, Q542), Critical activity (Y750) and Additional binding (M682) hotspot residues are highlighted with red circle. C-terminal domain of MLH1 (MLH1-C) is colored in green and C-terminal domain of PMS2 (PMS2-C) is colored in brown. (D) Dot graph of interface distance (log10 distance) by AlphaFold 3 competition test and interaction of pAE by RFdiffusion of MC (left) and MCA (right) binding proteins. Histogram of interface distance is above. Red line at 9.0157 Å indicates the original interface distance of the dimer complex of MLH1-C and PMS2-C. Yellow line at 9.7724 Å indicates our threshold value. MC, MCA and failed-MCA binders are highlighted by orange, green and blue box, respectively. (E-F) Fold-change in prime editing efficiency for *HEK3* CTT insertion (E) and *RNF2* TAC insertion (F) using designated MLH1 small binders, compared to PE2 efficiency. A 2.5-fold increase is highlighted with a red dotted line. MLH1 small binders that demonstrate more than a 2.5-fold increase in prime editing efficiency are marked with red dots. Bars represent mean values, and error bars represent the S.D. of n = 3 independent biological replicates.

To filter out unfavorable candidates, we used the predicted local distance difference test (pLDDT), averaged predicted aligned error of interchain residue pairs (interaction pAE), and root mean square deviation (RMSD) parameters from the RFdiffusion validation outputs (**Supplementary Tables 1-3**) ^23^. pLDDT indicates the stability of the binder protein; interaction pAE indicates the degree of binding affinity; and RMSD indicates design credibility. It has been shown that among these filtering parameters, interaction pAE is the most effective filtering parameter for binder design ^33^. Although interaction pAE characterizes the overall interaction between the binder and target well, it remains unclear how well the binder inhibits the binding of the original binding partner to the target protein. To address this problem, we further developed an AlphaFold 3 competition test, as following steps: i) we added a binder sequence to the existing dimer complex sequences and predicted the outcome of the trimer complex, ii) we selected five representative residue pairs around the interface between MLH1 and PMS2 to calculate the average atom-pair distance of the Cɑ atoms (**Fig. S1C**), and iii) using a histogram of the average distances, a filtering threshold was determined to exclude complexes that were unaffected by the binder (**Figs. 1D, S1B** and **S1D**). After conducting the AlphaFold 3 competition test, we identified three different groups in the distance histograms: the “first peak” indicating binders that could not induce an effective disturbance to the original complex, the “second peak” indicating binders that could partially bind and induce a small increase in the interface distance, and the “broad distribution” indicating binders that could bind fully and induce a complete replacement of the original binding partner, PMS2. We set the threshold to 9.7724 Å, as this only excludes the first unaffected cases. Resultantly, 20 MC binders and 23 MCA binders that passed the AlplaFold 3 competitions test were selected, respectively. For comparison, 17 MCA binders which failed the AlphaFold 3 competition test were named failed-MCA (**Figs. S2** and **S3**). Interestingly, we observed no significant structural differences between the MCA and failed-MCA binders (**Fig. S2B** and **S2C**), indicating that the AlphaFold 3 competition test appears to use latent knowledge about the protein complexes rather than their simple structural properties.

### AI-generated MLH1-SBs improve the PE efficiency

To evaluate the effect of the RFdiffusion-generated MLH1 small binders (i.e., MLH1-SBs) in the PE efficiency, we transfected each MLH1-SB along with PE2 ^7^ to HeLa cells **(Figs. 1E**, **1F**, and **S4)**. Consistent with our assumption, the addition of the “A” site for binder design showed a significant increase in the success rate, from 2/20 (10% for MC binders) to 12/23 (52% for MCA binders) in *HEK3* site and from 2/20 (10% for MC binders) to 3/23 (13% for MCA binders) in *RNF2* site (**Figs. 1E** and **1F**). We interpreted that the higher success rate of MCA binders over MC binders is caused by the effective local concentration of the binders at target interfaces through the non-competitive additional binding site. In contrast to the high success rate of the MCA binders, none of the failed-MCA binders (0/17 (0%)) enhanced the PE2 efficiency more than 2.5 fold, which demonstrates the clear discrimination ability of our AlphaFold 3 competition test. These failed-MCA binders were not recognized by the other filtering parameters, such as pLDDT, RMSD, and interaction pAE. This suggests that AlphaFold 3 successfully predicts binder disturbance of the original complex. Because the bare dimer complex between the binder and target lacks information regarding the competition between the binder and the original binding partner, trimer complex prediction is advantageous for identifying better candidates. Additionally, no clear correlation was observed between the fold change in PE efficiency and the AlphaFold 3 competition test scores, indicating the pass or fail of the test is more important (**Fig. S5**). For following experiments, we selected a 23th MCA binder, which was one of the five MLH1-SBs that increased the PE efficiency more than 2.5-fold in both *RNF2* and *HEK3* sites (**Figs. 1E** and **1F**) and was the smallest among the MCA binders (**Supplementary Table 2**).

### MLH1-SB is compatible with the La protein of PE7

A recent study reported the protection of the 3′ end of pegRNA with the La protein significantly increased the PE efficiency, generating PE7 architecture ^18^. As RNA protection and MMR are independent processes in cells, we hypothesized that the La protein is compatible with MMR inhibition ^13,18^. To examine this, we co-delivered MLH1dn or MLH1-SB with PEmax ^12^ (more optimized version of PE2) and PE7 ^18^ in HeLa cells, respectively (**Fig. 2A**). Through the inhibition of MMR, the overexpression of either MLH1dn or MLH1-SB significantly increased the insertion (*HEK3* CTT and *RNF2* TAC insertion), deletion (three-base pair (bp) deletion at *HEK3* and *RNF2*), and substitution (*HEK3* T to A, *HEK4* G to T and *RNF2* C to G) efficiencies with PEmax and PE7 (**Figs. 2B-2D**). Notably, MLH1-SB enhanced PE efficiency more than MLH1dn. Specifically, the average fold increase in PE efficiency for PEmax was 6.8-fold with MLH1dn and 18.1-fold with MLH1-SB (**Fig. 2E**), indicating that MLH1-SB showed 2.7-fold better activity than MLH1dn. Similarly, for PE7, the PE efficiency were increased by 2.7-fold with MLH1dn and by 9.6-fold with MLH1-SB (**Fig. 2E**), indicating that MLH1-SB showed 3.6-fold better activity than MLH1dn. These results indicate that the strategy for inhibition of MMR is compatible with the protection of the 3′ end of pegRNA and co-expression of MLH1-SB with PE7 exhibited the best PE performance.

**Figure 2.**
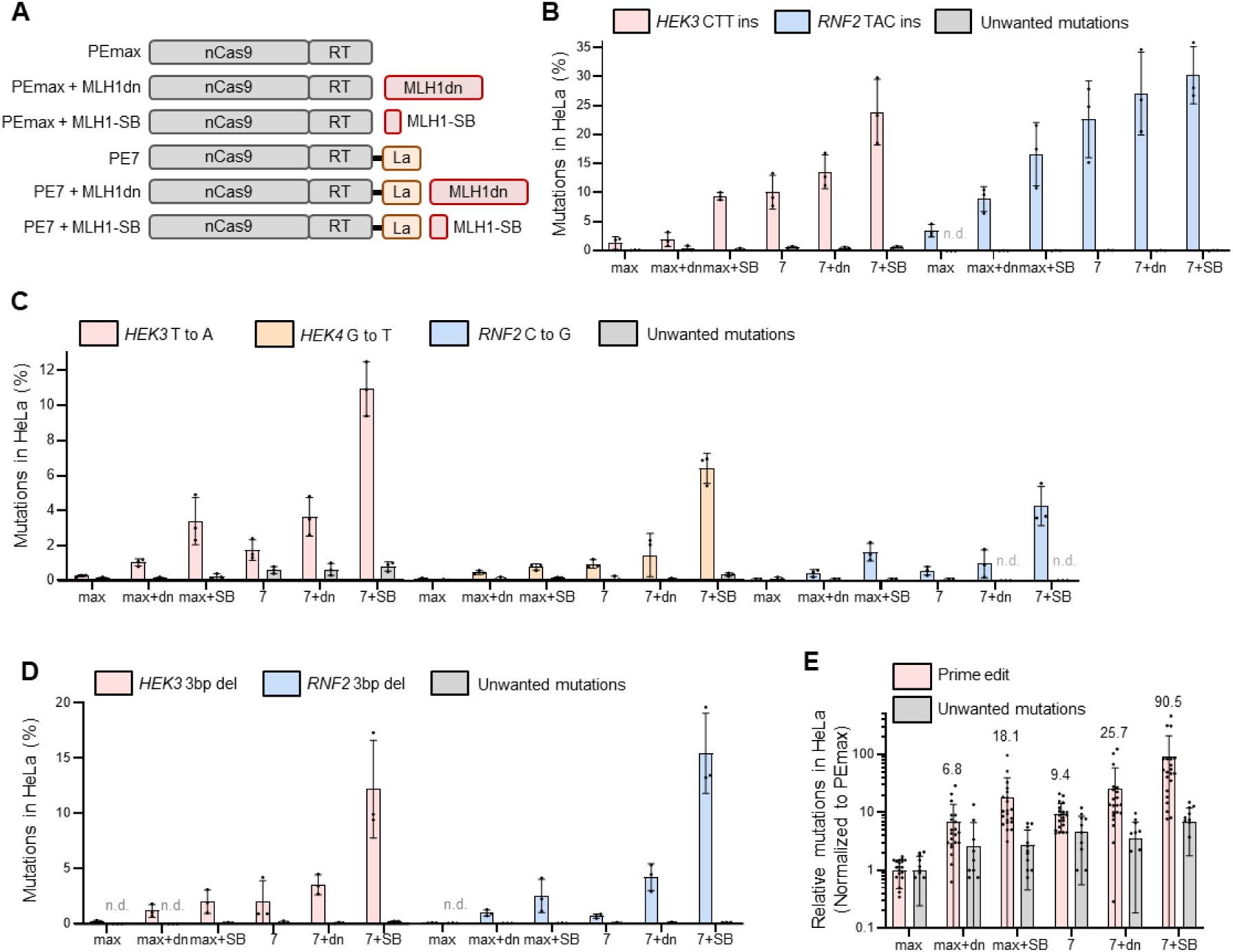
MLH1-SB is compatible with the La protein of PE7. (A) Schematics of prime editor architectures. nCas9, SpCas9 H840A nickase; RT, human codon-optimized MMLV-RT; MLH1dn, dominant negative MLH1; MLH1-SB, MCA-23 MLH1 small binder; La, small RNA-binding exonuclease protection factor La (B-D) Prime editing efficiency and unwanted mutations at designated loci and mutation for insertion (B) substitution (C) and deletion (D) mutation by PEmax and PE7 with control vector, MLH1dn or MLH1-SB expression vector in HeLa cells. Bars represent mean values, and error bars represent the S.D. of n = 3 independent biological replicates. n.d. for not detected data (E) Normalized prime editing and unwanted mutations by PEmax and PE7 with a control vector, MLH1dn, or MLH1-SB expression vector in HeLa cells. Experiments where unwanted mutations were not detected by PEmax were excluded from normalization. Numbers above the prime edits represent mean fold-change values. Bars represents mean values, and error bars indicate S.D. for n = 24 and n = 12 independent biological replicates, respectively, for prime edits and unwanted mutations.

### MLH1-SB binds to MLH1 and induces limited alteration in human cells

To investigate molecular mechanism of the selected MLH1-SB on enhancing PE efficiency in cells, we prepared two plasmids expressing MLH1-SB linked to an HA tag, named MLH1-SB-HA, and MLH1 linked to a 3 × FLAG tag, named the FLAG-MLH1. First, we conducted an immunoprecipitation assay using anti-HA beads, anti-HA antibody and anti-FLAG antibody (**Fig. 3A**). For HEK293T cells that lack MLH1 (**Fig. S6A**), we transfected the MLH1-SB-HA plasmid solely without the FLAG-MLH1 plasmid and observed no FLAG band (**Fig. 3A**). In contrast, when both MLH1-SB and the FLAG-MLH1 plasmids were co-transfected into HEK293T cells, the FLAG band was detected, indicating that MLH1-SB directly binds to MLH1 in cells. Next, we analyzed the localization of MLH1-SB in relation to the location of MLH1 using immunofluorescence (IF) staining (**Fig. 3B**). After transfecting an empty vector (“Mock”), the MLH1-SB-HA plasmid, the FLAG-MLH1 plasmid, or both of MLH1-SB-HA and FLAG-MLH1 plasmids into HEK293T cells, we stained the cells with Hoechst (blue), anti-FLAG antibody (magenta), and anti-HA antibody (green). Results showed that FLAG staining predominantly occurred in the nucleus demonstrating the typical nuclear localization of MLH1. Conversely, HA staining was primarily observed in the cytoplasm, suggesting that MLH1-SB is predominantly located in the cytoplasm. Interestingly, we observed increased green fluorescence signals in the nucleus of both MLH1- and MLH1-SB-expressing cells. Given that MLH1-SB does not involve nuclear localization signal (NLS) peptides, these results indicate that MLH1-SB binds to MLH1 containing NLS peptides and co-transports into the nucleus with the MLH1. To further confirm whether the increase of PE efficiency by MLH1-SB depends on MMR in cells, we transfected PEmax and PE7 along with MLH1dn or MLH1-SB into MMR-deficient HEK293T cells (**Figs. 3C-3F**). Intriguingly, the PE efficiency of both PEmax and PE7 remained comparable under the expression of MLH1dn and MLH1-SB (**Fig. 3F**), suggesting that the PE enhancement by MLH1-SB is MMR-dependent.

**Figure 3.**
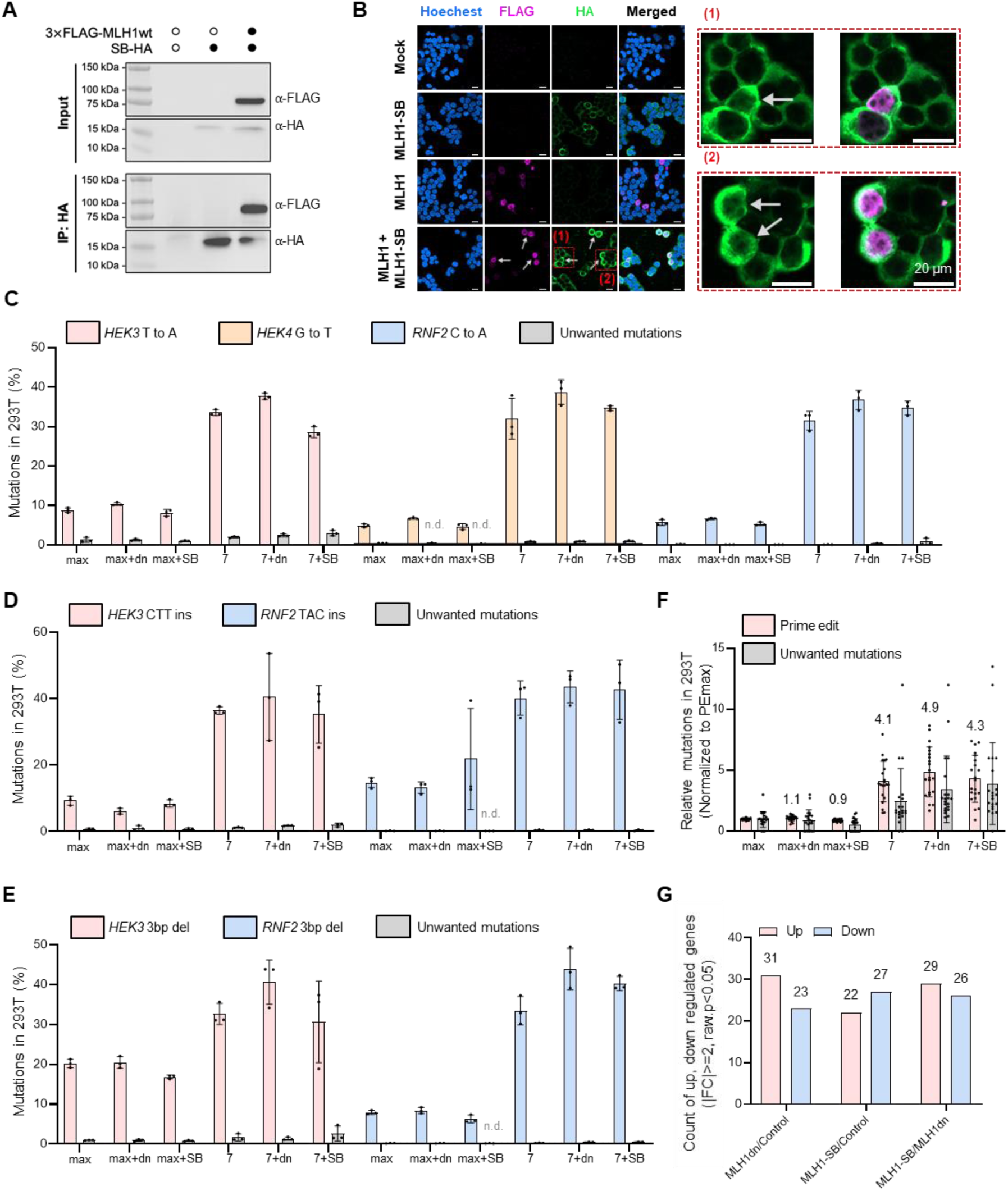
MLH1-SB binds to MLH1 and induces limited alteration in human cells. (A) Blot of immunoprecipitation of HEK293T cells transfected with MLH1-SB with HA tag only or MLH1-SB with HA tag and wild-type MLH1 with 3 x FLAG tag. (B) Immunofluorescence image of HEK293T cells transfected with different constructs. Cells were stained with Hoechst (blue) to visualize nuclei, anti-Flag antibody (magenta) to detect FLAG-tagged MLH1, and anti-HA antibody (green) to detect HA-tagged MLH1-SB. (Left) Mock, Cells transfected with an empty vector, showing no specific staining for Flag or HA; MLH1-SB, Cells expressing MLH1-SB show HA staining in the cytoplasm; MLH1, cells expressing MLH1 show Flag staining primarily in the nucleus; MLH1 + MLH1-SB, MLH1-SB in cells co-transfected with MLH1 and MLH1-SB display both nuclear and cytoplasmic localization, indicating potential interaction or co-localization. (Right) Higher magnification images of the regions marked in the merged images of the MLH1 + MLH1-SB. (1), Enlargement of the indicated region showing both nuclear and cytoplasmic signals; (2), Another magnified region showing similar co-localization. Arrows indicating cells with high MLH1 expression. Scale bars represent 20 µm. (C-E) Prime editing efficiency and unwanted mutations at designated loci and mutation for substitution (C) insertion (D) and deletion (E) mutation by PEmax (max) and PE7 (7) with control vector, MLH1dn (+dn) or MLH1-SB (+SB) expression vector in HEK293T cells. Bars represent mean values, and error bars represent the S.D. of n = 3 independent biological replicates. n.d. for not detected data (F) Normalized prime editing and unwanted mutations by PEmax and PE7 with a control vector, MLH1dn, or MLH1-SB expression vector in 293T cells. Numbers above the prime edits represent mean fold-change values. Bars represents mean values, and error bars indicate S.D. for n = 24 independent biological replicates. (G) Bar graph summarizing the number of upregulated and downregulated genes (|Log2FC| ≥ 2, raw p < 0.05) in each designated condition. Numbers above the bars represent the number of differentially expressed genes.

We further evaluated the effect of the transient expression of MLH1dn and MLH1-SB on the cell’s characteristics through a transcriptome-analysis. This was followed by the transfection of a control vector (PuroR expression vector), MLH1dn, or MLH1-SB into human induced pluripotent stem cells (hiPSCs). Of note, 49–55 differentially expressed genes (DEGs) were observed among the three different transcriptomes of hiPSCs (**Figs. 3G** and **S6B**). Gene ontology enrichment analysis ^34^ of the DEGs indicated that the biological characteristics of the cells were comparable (**Figs. S6C-S6E**). Additionally, the viability of the HeLa cells transfected with MLH1dn or MLH1-SB were comparable with the cells transfected with control vector (**Fig. S6F**). Taken together, we concluded that MLH1-SB expression does not induce serious alterations in human cells.

### MLH1-SB fused prime editors exhibit enhanced PE efficiency

Given that the co-expression of MLH1-SB significantly increases the PE efficiency, we attempted to incorporate MLH1-SB into the PE architectures for the establishment of improved prime editors. For this purpose, we connected MLH1dn, MLH1-SB, and MLH1-SB with a NLS (referred to as MLH1-SB-NLS), or both MLH1-SB-NLS and MLH1-SB (referred to as MLH1-SB2), to PE7 via a 2A peptide which is an oligopeptide that facilitates the cleavage of proteins during translation (**Fig. 4A**). Consequently, we named PE7 connected through the 2A system with MLH1-SB as PE7-SB, PE7 with MLH1-SB-NLS as PE7-SB-NLS, and PE7 with MLH1-SB2, as PE7-SB2. In addition, we fused MLH1-SB to either the N-terminal or C-terminal end of PE7 using an XTEN linker (Xt), named PE7-N-SB and PE7-C-SB, respectively. We delivered these prime editors to HeLa cells for PE in three different target (*HEK4* G to T and 3 bp deletion in *HEK3* and *RNF2*). Among the prime editors that we tested, PE7-SB2 was comparable to PE7-SB and PE7-SB-NLS but exhibited slightly higher PE efficiency than others (**Figs. 4B** and **S7**). Interestingly, PE7 linked to MLH1dn (named PE7-MLH1dn) did not show significant increased PE efficiency over PE7 in average, in contrast to the PE7-SB platforms, interpreting that the small size (82 amino acids) of MLH1-SB allowed it to be integrated into prime editor.

**Figure 4.**
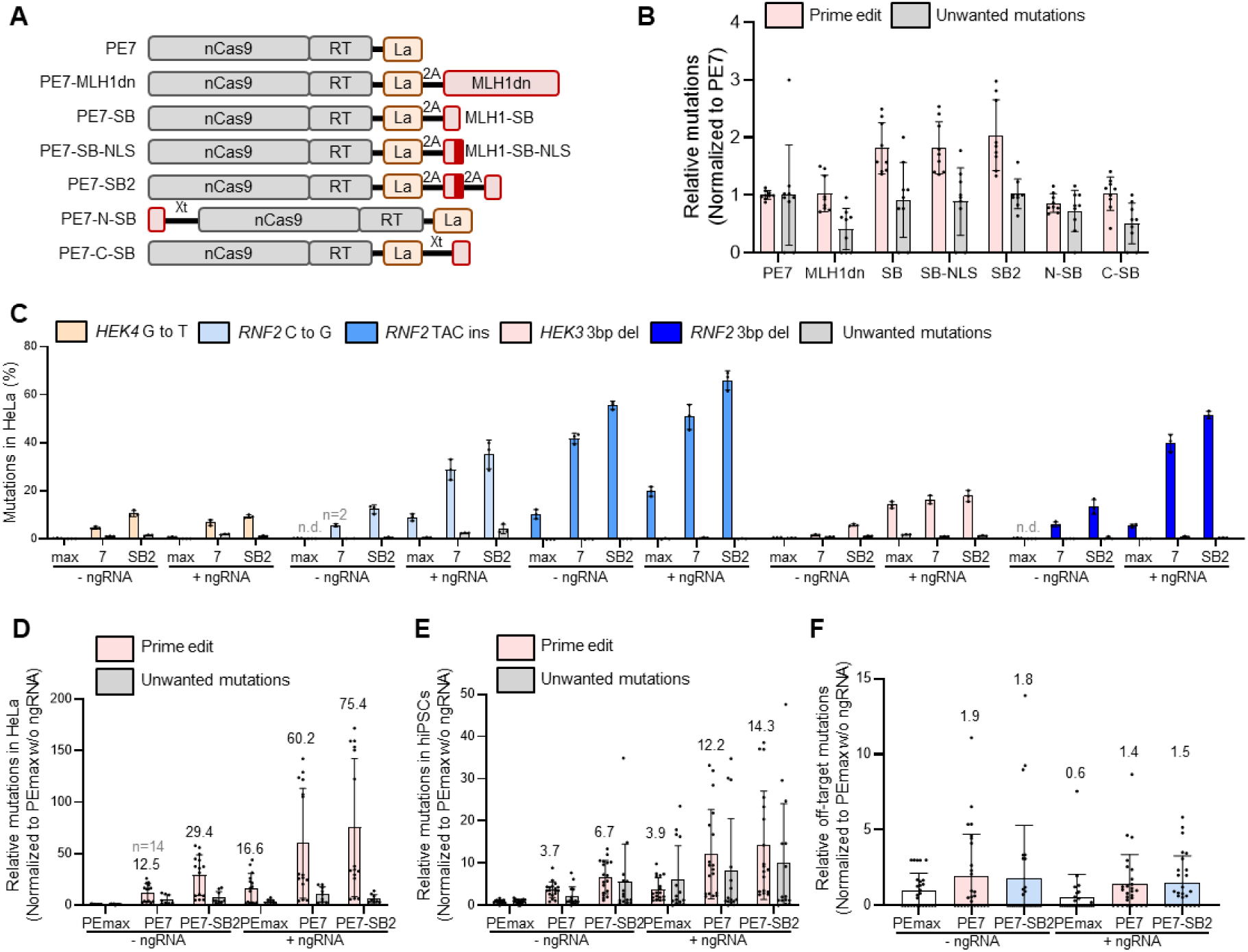
MLH1-SB fused prime editors exhibit enhanced PE efficiency. (A) Schematics of prime editor architectures. nCas9, SpCas9 H840A nickase; RT, human codon-optimized MMLV-RT; MLH1dn, dominant negative MLH1; MLH1-SB, MCA-23 MLH1 small binder; MLH1-SB-NLS, MLH1-SB with nucleus localization sequence (NLS); La, small RNA-binding exonuclease protection factor La; 2A, 2A oligopeptide; Xt, SGGS-XTEN16-SGGS linker. (B) Normalized prime editing and unwanted mutations by PE7, PE7 with MLH1dn (MLH1dn), PE7 with MLH1-SB (SB), PE7 with MLH1-SB-NLS (SB-NLS), PE7 with both MLH1-SB and MLH1-SB-NLS (SB2), PE7 linked to MLH1-SB at N-terminal (N-SB) or C-terminal (C-SB) of PE7 in HeLa cells. Numbers above the prime edits represent mean fold-change values. Bars represents mean values, and error bars indicate S.D. for n = 9 independent biological replicates. (C) Prime editing efficiency and unwanted mutations at designated loci and mutation by PEmax (max), PE7 (7), and PE7-SB2 (SB2) with or without nicking gRNA (ngRNA) in HeLa cells. Bars represent mean values, and error bars represent the S.D. of n = 3 independent biological replicates except the designated replicates. n.d. for not detected data. (D) Normalized prime editing and unwanted mutations by PEmax, PE7, and PE7-SB2 with or without ngRNA in HeLa cells. Experiments where unwanted mutations were not detected by PE7 were excluded from normalization. Bars represents mean values, and error bars indicate S.D. for n = 15 and n = 9 independent biological replicates, respectively, for prime edits and unwanted mutations except the designated replicates. (E) Normalized prime editing and unwanted mutations by PEmax, PE7, and PE-SB2 with or without ngRNA in hiPSCs. Bars represents mean values, and error bars indicate S.D. for n = 18 and n = 15 independent biological replicates, respectively, for prime edits and unwanted mutations. (F) Normalized off-target editing by PEmax, PE7, and PE-SB2 with or without ngRNA in HeLa cells. Bars represents mean values, and error bars indicate S.D. for n = 27 independent biological replicates.

We further evaluated whether PE7-SB2 was also compatible with addition of ngRNA. To achieve this, we transfected PEmax, PE7, and PE7-SB2 with or without ngRNA for substitution (*HEK4* G to T and *RNF2* C to G), insertion (*RNF2* TAC insertion) and deletion (3-bp deletion in *HEK3* and *RNF2*) mutations in HeLa cells (**Fig. 4C**). As proven in above experiments, for the various cases, PE7-SB2 improved PE efficiency by 29.4-fold compared to PEmax and by 2.4-fold compared to PE7 (**Fig. 4D**). Moreover, PE7-SB2 with the addition of ngRNA further improved PE efficiency by 2.6-fold compared to the single PE7-SB2, implying that PE7-SB2 is compatible with addition of ngRNA. The ngRNA addition effect was also assessed in hiPSCs for substitution (*HEK3* T to A and *HEK4* G to T), insertion (*HEK3* CTT insertion and *RNF2* TAC insertion) and deletion (3-bp deletion in *HEK3* and *RNF2*) (**Figs. 4E** and **S8**). Similar to the results in HeLa cells, for the various cases, PE7-SB2 showed 6.7-fold and 1.8-fold increases in PE efficiency compared to PEmax and PE7, respectively in hiPSCs. The addition of ngRNA to PE7-SB2 further improved the PE efficiency by 2.1-fold compared to PE7-SB2 without ngRNA.

Next, we examined the editing efficiency at well-defined off-target sites: three off-target sites for *HEK3* and *HEK4* predicted by CIRCLE-seq ^12,35^, and five off-target sites for *RNF2* predicted by Cas-OFFinder ^36^ (**Fig. S9**). After transfecting PEmax, PE7, or PE7-SB2 with each pegRNA targeting *HEK3*, *HEK4* and *RNF2* site in HeLa cells, each target site was amplified and subject to targeted deep sequencing. Results showed that the average off-target editing of PE7-SB2 was higher than PEmax but was comparable with PE7, indicating that MLH1-SB2 does not generate significant off-target editing additionally (**Figs. 4F** and **S9**).

## Discussion

The rapid development of generative AI tools is changing the research environment not only in academia but also in industry. In this study, using RFdiffusion ^23^ and AlphaFold 3 ^20^, we generated small proteins ranged from 50 to 150 amino acids in size, which could bind to the dimeric interface of MLH1 and PMS2, to inhibit MMR, thereby enhancing the PE efficiency. Among the 60 generated MLH1-SBs, we selected the 23th MLH1-SB with a size of 82 amino acids and PE efficiency that increased 2.8-fold for *RNF2* TAC insertion and 4.4-fold for *HEK3* CTT insertion. In contrast to MLH1dn, which is 753 amino acids in size, the MLH1-SB’s small size allowed it to be integrated into pre-existing PE architectures via the 2A system, creating a novel PE-SB platform. It is noteworthy that the increase of PE efficiency by MLH1-SB was greater than that by MLH1dn, a widely used protein for MMR inhibition ^12^. Moreover, MLH1-SB was also compatible with La protein used in PE7 ^18^ and/or the addition of nicking sgRNA used in PE3 or PE5. By integrating MLH1-SB into PE7, we have successfully developed an improved PE architecture called PE7-SB, which demonstrates the highest PE efficiency to date (29.4-fold over PEmax and 2.4-fold over PE7 in HeLa cells). Considering the payload capacity of delivery methods for *in vivo* gene therapy such as adeno-associated virus (AAV) ^37^ and lipid nanoparticles (LNPs) ^38^, MLH1-SB that significantly enhances PE while maintaining minimal size is expected to facilitate the development of highly effective *in vivo* gene therapy.

To the best of our knowledge, this study represents the first instance of using AI technologies for target pathway inhibition to enhance genome editing efficiency. Until now, the application of RFdiffusion has primarily been limited to enhancing binding affinity ^23,27^ and designing de novo functional proteins ^39,40^. But, we here demonstrated that AI-generated SBs that target the dimeric interface of protein of interest (POI) can successfully inhibit the function of the target protein complex. This finding expands the application of RFdiffusion and the toolbox for the transition inhibition of the POI. We expect that this approach will prompt breakthroughs in diverse biological and biomedical research areas.

In addition, we present guidelines for generating AI-based SBs to inhibit the formation of protein complexes. We generated MLH1-SBs via RFdiffusion under two conditions: MC and MCA binding. MC binding conditions produced compact, sphere-like proteins, whereas MCA binding conditions resulted in rod-like proteins that featured a long alpha chain. Notably, only two out of the 20 MC binders increased PE efficiency by more than 2.5-fold, while 12 out of the 23 MCA binders achieved a greater than 2.5-fold increase in PE efficiency. These findings suggest that a broader interaction surface between the target protein and small binders substantially contributes to effective inhibition. Nevertheless, the fourth MC binder, despite being only 51 amino acids in length and exhibiting a 2.8 to 4-fold increase in PE efficiency, contradicts this pattern. This indicates that a wide interacting surface is beneficial but is not an absolute requirement for strong inhibition of dimer formation. For the assessment of AI-generated MLH1-SBs, RFdiffusion provides assessment tools of RMSD, interaction pAE, and pLDDT ^23^. Although these parameters predict protein folding and binding stability, they provide limited information about the protein’s competitive efficacy with other binding partners. To address this limitation, we implemented the AlphaFold 3 competition test. In this test, the original complex proteins (e.g., MLH1 C-terminal and PMS2 C-terminal) were folded together with a small binder using AlphaFold 3 ^20^. A shift in the distance between the original complex proteins indicates the competitive ability of the small binder. Intriguingly, failed-MCA binders, which were generated under MCA conditions but failed the AlphaFold 3 competition test, did not increase prime editing efficiency by greater than 2.5-fold. This suggests that the AlphaFold 3 competition test was effective at predicting the competitive efficacy of small binders. However, the fold change in PE efficiency did not correlate with the AlphaFold 3 competition test scores, indicating that an overall pass or fail of the test was more critical than the absolute value of the score.

Furthermore, it is notable that we could quickly generate MLH1-SBs without local high computing power using available web-based AI tools (by Colab version RFdiffusion, three days) and effectively filter the candidates (by AlphaFold 3 binding competition, one day). High success rate due to our unique approaches, including multi-utility of hotspots and AlphaFold 3 competition filtering, also saves time when performing the trials. Our pipeline has the potential to be an efficient way to develop a SB for many purposes. Recent advances regarding AI-based protein models, including RoseTTAFold All-Atom ^22^, will expand the ability of protein generation algorithms and enable us to design binders to more general targets, such as ligands and nucleic acids. A more accurate protein complex folding model will also improve the competitive filtering method.

Although our study successfully established improved PE platforms and provided valuable insights into the integration of AI technologies with genome editing, there are a few limitations to be addressed further. In this study, we inhibited the MMR pathway to enhance PE activity. However, the loss of MMR is closely related to the accumulation of genome-wide mutations and the development of cancer ^41–43^. A recent study revealed that the transient expression of MLH1dn in mouse embryos increased the number of genome-wide small deletion mutations ^44^. Thus, further investigation of MLH1-SB-mediated effects is required. Also, we generated SBs to disrupting the formation of the MutLα complex, but it is ambiguous whether our approach can be applicable for proteins that operate solely. Thus, the potential of SBs to inhibit the function of monomeric proteins requires further evaluation. Moreover, in addition to inhibition, it is necessary to develop SBs capable of activating specific pathways to enhance genome editing.

Previously, we and others found that the enhancement of PE efficiency through the depletion of MMR was limited to edits up to approximately 10 bps in length ^14,45^. Consistently, the MMR pathway is known to be regulated to a length of approximately 13 bps ^13,46^. Given the importance of long-sequence editing by prime editors (e.g., insertion of AttB and AttP sequences for targeted integration)^47,48^, exploring and manipulating the cellular mechanisms that influence PE efficiency of long-sequences may serve as a foundation for future research. Starting with this study, generative AI will provide significant benefits and insights for upgrading and developing new genome editing tools beyond the prime editor.

## Supporting information

Supplementary informations

## Acknowledgements

We thank Prof. David R Liu for his critical comments. Most analysis of sequencing data was carried out using the computing server at the Genomic Medicine Institute Research Service Center. This research was supported by grants from the National Research Foundation of Korea (NRF) No. 2021M3A9H3015389, No. RS-2024-00451880, and SRC - NRF2022R1A5A102641311 to S.B., and by the Korean Fund for Regenerative Medicine (KFRM) grant No. RS-2024-00332601 to S.B.

## Author contributions

J.-C.P., H.U., and S.B. conceived this project; H.U. developed bioinformatics algorithms; J.-C.P., Y.-W.K., and Y.E.O. performed cell experiments; S.B. supervised this project; J.-C.P., H.U., and S.B. wrote the manuscript with the help of all other authors.

## Additional information

Supplementary Information accompanying this paper is available at http://

## Materials and Methods

### Small binding protein generation with RFdiffusion

A customized code modified from the original Colab version of RFdiffusion integrated with ProteinMPNN and Alphafold 2 was used for our binder generation. The dimeric structure of MLH1-C (486-751) and PMS2-C (606-862) was generated by Alphafold 3 via the official website. Using the generated MLH1-C structure as a target protein, we specified 3 hotspots, W538, Q542 and Y750, for MC binders, and added one more hotspot, M682, for MCA binders. Most of the adjustable parameters were set to default values for binder design as recommended by the RFdiffusion developers. Some important options are here. In the RFdiffusion subsection, iterations, 50; in the ProteinMPNN subsection, num_seqs, 2; mpnn_sampling_temp, 0.0001; rm_aa, ‘C’; and in the AlphaFold 2 subsection, initial_guess, ‘TRUE’; num_recycles, 3; use_multimer, ‘TRUE’ as ‘50-150’ for MC binders and as ‘70-150’ for MCA binders. In the Colab environment, A100 GPU was used for our runtime. Total running time was about 20 hours using 6 Colab-runtimes simultaneously. The cost was normally 0.01∼0.1$ and 1∼10min per 1 candidate in 1 Colab-runtime. Due to occasional runtime aborts which is unavoidable for Colab environment, actual consumed time was 3 days for the binder generations. 1978 MC binders and 821 MCA binders were generated until we got enough filtered candidates. Filtering was done by threshold values, 7.5 for interaction pAE, 0.85 for pLDDT, 1.5 Å for RMSD. Remaining candidates were used for AlphaFold 3 competition test.

### AlphaFold 3 competition test

The sequences of the selected candidates were used for structure prediction of trimer complex with MLH1-C and PMS2-C by Alphafold 3 via the official website. The average atom-pair distance of the Cɑ atoms of the interface between MLH1-C and PMS2-C (i.e., interface distance) was calculated using five selected residue-pairs, L540, L743, Q542, C756, N739 in MLH1-C and I688, I853, N683, E705, Q861 in PMS2-C. For comparison, the interface distance in the dimeric structure of MLH1-C and PMS2-C was also calculated, as 9.0157 Å. Using a threshold value, 9.7724 Å, if the interface distance of the trimer structure was larger than it, the candidate was passed. The passed candidates were ranked in the interaction pAE values. The top ranks were selected for experimental validation. The final numbers were 20 for MC, 23 for MCA, and 17 for failed-MCA.

### Plasmid construction

The proteins sequences of MLH1-SBs formed by RFdiffusion were transformed into genetic sequence by utilizing GenSmart Optimization. The codon-optimized MLH1-SBs sequences were synthesized by Gene Fragments from TwistBioscience. The gene fragments of MLH1-SBs were amplified by PCR and integrated into PCR-linearized CMV-PuroR-2A backbone vector via Gibson Assembly. To generate MLH1dn expression vector, we amplified MLH1dn sequence from pCMV-PE4stem plasmid (addgene, 208768). The amplified MLH1dn is integrated into PCR-linearized CMV-2A-Puro backbone vector with Gibson Assembly. To generate pCMV-PE7max, cellular RNA were extracted from HEK293T cells using TRIZol-Chloroform. 1 µg of RNA were reverse trancribed using ReverTra Ace® qPCR RT Master Mix (TOYOBO, FSK-101F) according to manufacturer’s protocol. La(1-194); also known ad ‘SSB’ cDNA were amplified using KOD -Plus-Neo (TOYOBO, TOKOD-401) and cloned into pCMV-PEmax (addgene, 174820) vectors using BsrGI-HF (NEB, R3575S) and SapI (NEB, R0569S) with SGGSx2-XTEN16-SGGSx2 linker. To generate MLH1-SB-NLS sequence, codon of MLH1-SB is re-optimized with GenSmart Optimization. The re-optimized MLH1-SB with NLS sequence is synthesized by Gene Synthesis of BIONICS and amplified with PCR for further cloning. The MLH1-SB-NLS sequence was codon re-optimized by GenSmart Optimization and synthesized by BIONICS and subsequently cloned into the PE7max vector using KpnI-HF (NEB, R3142L) and AgeI-HF (NEB, R3552L) restriction enzymes. To generate PE7-MLH1-SB-NLS, site-directed mutagenesis was performed using PCR with Platinum™ SuperFi II DNA Polymerase (Invitrogen, 361250). Codon re-optimized MLH1-SB-NLS assembled into the PE7-MLH1-SB construct through Gibson Assembly for the generation of PE7 with both MLH1-SB and MLH1-SB-NLS. MLH1-SB is integrated into 5’ or 3’ end of PE7 vector which is linearized with PCR to have SGGS-XTEN16-SGGS at the 5’ or 3’ end of PE7 via Gibson Assembly for the generation of PE7-N-term-SB and PE7-C-term-SB vector respectively. PCR was performed with KOD Multi & Epi PCR kit (TOYOBO) and Gibson Assembly was performed with Gibson Assembly Master Mix (NEB, E2611).

### Cell line culture condition

HEK293T (ATCC, CRL-11268), Hela (ATCC, CLL-2) and U2OS (ATCC, HTB-96) cells were cultivated in DMEM media (Welgene, LM001-05) containing 10% FBS (Welgene, PK004) and 1% Antibiotics (Welgene, LS203-1) in a humidified incubator at 37 °C with 5% CO2. For transfer of cells, cells were washed with DPBS (Welgene, LB001-01) and detached with Trypsin-EDTA 0.25% (Welgene, LS015). After detachment of cells, DPBS with 10% FBS was added for the inactivation of Trypsin-EDTA. Cells were counted and seeded to cell culture plate with cell culture media. hiPSCs (CMC-hiPSC-022, Catholic University of Korea) were grown in Essential 8™ media (ThermoScientific, A1517001) on cell culture plates coated with iMatrix-511 (REPROCELL, NP892-012). For transfer of hiPSCs, cells were washed with DPBS and detached with ACCUTASE™ (Stem Cell Technologies, 07922). Detached cells were washed with DPBS for two times. Washed cells were seeded to iMatrix-511 coated cell culture plate containing Essential 8™ media. To prevent dissociated cell death of hiPSCs, 10 μM of ROCK-inhobitor Y27632 dihydrochloride (MCE, HY-10583) is added to the cell culture media when transferred. hiPSCs were incubated at 37 °C with 5% CO2.

### Transfection and genomic DNA extraction

For transfection of plasmid DNAs, 293T, U2OS and HeLa cells were washed with DPBS (Welgene, LB001-01) and detached with Trypsin-EDTA 0.25% (Welgene, LS015) and washed with DMEM with 10% FBS. Washed cells were counted and seeded into 48-well cell culture plate (SPL, 30048) with 5 × 10^4^ cells were per well. In case of MLH1-SB test experiment (Figs. 1e,f), 2.5 × 10^4^ of HeLa cells were seeded into 96-well cell culture plate. After 24 hours from seeding, media was replaced with fresh DMEM with 10% FBS. For MLH1-SB test experiment (Figs. 1e,f), 60 ng of pCMV-PE2 vector (Addgene, 132775), 20 ng of pegRNA vector and 20 ng of MLH1-SB expression vector was mixed with 0.5 μl of Lipofectamin 2000 (ThermoFisher, 11668027) followed by the manufacturer’s instruction and treated to HeLa cells. For PE2 or PE7 with MLH1dn or MLH1-SB experiment (Fig. 2), 120 ng of pCMV-PEmax vector (Addgene, 174820) or PE7 vector, 40 ng of pegRNA vector and 100 ng of control vector (CMV-Puro^R^ vector), MLH1dn or MLH1-SB expression vector were mixed with 1 μl of Lipofectamin 2000 and treated to HeLa and 293T cells. For PE8 screening test experiment (Fig. 3a), 180 ng of each form of prime editor (i.e., PE7, PE7-dn, PE7-SB, PE7-SB-NLS, PE7-2xSB, PE7-N-term-SB and PE7-C-term-SB) and 60 ng of pegRNA are mixed with 1 μl of Lipofectamin 2000 and treated to HeLa cells. For PE8 test experiment (Figs. 3c-h), 180 ng of pCMV-PEmax vector (Addgene, 174820), PE7 vector or PE7-2xSB vector, 60 ng of pegRNA vector and 20 ng of nicking gRNA (for + ngRNA condition) are mixed with 1 μl of Lipofectamin 2000 and added to HeLa and U2OS cells. For the transfection of hiPSCs, cells were detached with Accutase (Stem Cell Technologies, 07922) and washed with DPBS. Cells were counted and 3 × 10^4^ cells were seeded to 48-well cell culture plate. 180 ng of pCMV-PEmax vector (Addgene, 174820), PE7 vector or PE7-2xSB vector, 60 ng of pegRNA vector and 20 ng of nicking gRNA (for + ngRNA condition) are mixed with 1 μl of Lipofectamin Stem (ThermoFisher, STEM00015) followed by the the manufacturer’s instruction and added to iPSCs. After 72 hours from transfection, media was removed, and cells were washed with DPBS. For extraction of genomic DNA, washed cells were lysed with 25 μl (for 96 well experiment) or 60 μl (for 48 well experiment) of lysis buffer composed of 40 mM Tris, pH8.0, 1% Tween-20, 0.2 mM EDTA, 0.2% Nonidet P-40 and 2 mg/ml of protease K. The genomic DNA extract was incubated at 60 °C for 15 min and 95 °C for 5 min for activation and inactivation of Protease K.

### Co-immunoprecipitation

HeLa cells (0.5 million) were seeded into 6-well plates 12-16 hours before transfection. The next day, cells were transfected with 2 µg of plasmid DNA (either MLH1-SB-HA or a combination of MLH1-SB-HA and 3xFlag-MLH1) using Lipofectamine 2000 (Invitrogen #11668019) according to the manufacturer’s instructions. Following transfection, cells were incubated at 37°C with 5% CO₂ for 48 hours. Forty-eight hours post-transfection, the culture medium was removed, and the cells were washed once with 1X DPBS (Welgene, LB001-01). RIPA buffer, supplemented with cOmplete™, Mini Protease Inhibitor Cocktail (Roche, 11836153001), was added to the cells for lysis. The cell lysates were passed through a 26G syringe needle several times to ensure complete lysis and incubated on ice for 30 minutes, with vortexing every 10 minutes.

The lysates were incubated with pre-washed, equilibrated Pierce™ Anti-HA Magnetic Beads (Invitrogen, 88836) and rotated overnight at 4°C to facilitate binding. The following day, the beads were separated using a magnetic stand and washed twice with 0.05% TBS-T. The bound proteins were eluted by adding 2X sample buffer and heating at 95°C for 5 minutes. The eluted proteins were resolved on 4–20% Mini-PROTEAN® TGX™ Precast Protein Gels (Bio-Rad, 4561094) and transferred onto 0.45 µm PVDF membranes (Bio-Rad, 1620177). Detection of 3xFlag-MLH1 was performed using the DYKDDDDK Tag Monoclonal Antibody (Invitrogen, MA1-91878), while detection of MLH1-SB-HA was carried out using the HA Tag Monoclonal Antibody (Invitrogen, 26183).

### Targeted deep sequencing

Each on-target site was amplified through two rounds of PCR. Briefly, initial step PCR1 amplified the genome of the interest site via target site primers containing illumina Forward and reverse sequence adaptor. In PCR1, 1ul of genomic DNA extract was amplified using KOD-Multi & epi (TOYOBO, KME-101) and was performed with the amplify conditions: 94 °C for 2 min, 32 cycles of (98 °C for 10 s, 58 °C for 20 s, and 68 °C for 12s). The PCR1 product was used in PCR2 for Illumina sequencing. 1µl of PCR1 product and proceeded with Illumina sequence index primers. Amplify conditions: 94 °C for 2 min, 20 cycles of (98 °C for 10 s, 60 °C for 20 s, and 68 °C for 12s). For confirm the PCR product, 1.5% agarose gel electrophoresis. (Condalab, 8100) The PCR2 products were pooled and purified with PCR purification Kit (GeneAll, 103-150). The libraries were analyzed using the Illumina Miniseq instrument, and the miniseq result were analyzed by Cas-analyzer((http://www.rgenome.net/be-analyzer/).

### RT-qPCR

For extraction of total RNA, 293T, U2OS, iPSCs and HeLa cells were detached and washed with DPBS. Washed cells were lysed with 500 μl of TRIzol Reagent (Invitrogen, 15596018) and total RNA was extracted following the supplier’s instruction. 500 ng of RNA was mixed with 2 μl of PrimeScriptTM RT reagent kit (TaKaRa, RR037B) and distilled water (DW) up to 10 μl of total volume for cDNA synthesis. Mixed reagents were reacted at 37 °C for 15 min and 85 °C for 30 sec. Synthesized cDNA was diluted with 200 μl of DW. 10 μl of iTaq Universal SYBR Green Supermix (Bio-rad, 1725124), 0.5 μl of each PCR forward and reverse primer (20 pmol) and 9 μl of diluted cDNA was added to 96-well qPCR plate. qPCR was performed with CFX 96 Real-Time PCR Detection System (Bio-rad) with following cycling conditions: 95 °C for 3 min, and 50 cycles of (95 °C for 10 sec, 55 °C for 10 sec, 72 °C for 30 sec). Relative MLH1 expression level was calculated by using GAPDH as control.

### mRNA sequencing

Total RNA concentration was calculated by Quant-IT RiboGreen (Invitrogen, #R11490). To assess the integrity of the total RNA, samples are run on the TapeStation RNA screentape (Agilent, #5067-5576). Only high-quality RNA preparations, with RIN greater than 7.0, were used for RNA library construction. A library was independently prepared with 10 ng of total RNA for each sample by SMARTer ultra-low input RNA Sample Prep Kit (Clontech Laboratories, Inc., CA, USA). The first step in the workflow involves purifying the poly-A containing mRNA molecules using poly-T-attached magnetic beads. Following purification, the mRNA is fragmented into small pieces using divalent cations under elevated temperature. The cleaved RNA fragments are copied into first strand cDNA using SMARTerscribe reverse transcriptase (Clontech) and 3’ SMART CDS primers. This is followed by second strand cDNA synthesis using DNA Polymerase I, RNase H and dUTP. These cDNA fragments then go through an end repair process, the addition of a single ‘A’ base, and then ligation of the adapters. The products are then purified and enriched with PCR to create the final cDNA library.

The libraries were quantified using KAPA Library Quantificatoin kits for Illumina Sequecing platforms according to the qPCR Quantification Protocol Guide (KAPA BIOSYSTEMS, #KK4854) and qualified using the TapeStation D1000 ScreenTape (Agilent Technologies, # 5067-5582). Indexed libraries were then submitted to an Illumina NovaSeq (Illumina, Inc., San Diego, CA, USA), and the paired-end (2×100 bp) sequencing was performed by the Macrogen Incorporated.

### mRNA sequencing data

We preprocessed the raw reads from the sequencer to remove low quality and adapter sequence before analysis and aligned the processed reads to the Homo sapiens(hg38) using HISAT v2.1.0(1). HISAT utilizes two types of indexes for alignment (a global, whole-genome index and tens of thousands of small local indexes). These two types’ indexes are constructed using the same BWT (Burrows–Wheeler transform) a graph FM index (GFM) as Bowtie2. Because of its use of these efficient data structures and algorithms, HISAT generates spliced alignments several times faster than Bowtie and BWA widely used. The reference genome sequence of Homo sapiens(hg38) and annotation data were downloaded from the UCSC table browser (http://genome.uscs.edu). Transcript assembly and abundance estimation using StringTie(2, 3). After alignment, StringTie v2.1.3b was used to assemble aligned reads into transcripts and to estimate their abundance. It provides the relative abundance estimates as Read Count values of transcript and gene expressed in each sample.

### Statistical analysis of gene expression level

The relative abundances of gene were measured in Read Count using StringTie. We performed the statistical analysis to find differentially expressed genes using the estimates of abundances for each gene in samples. Genes with one more than zeroed Read Count values in the samples were excluded. To facilitate log2 transformation, 1 was added to each Read Count value of filtered genes. Filtered data were log2-transformed and subjected to RLE normalization. Statistical significance of the differential expression data was determined using nbinomWaldTest using DESeq2 and fold change in which the null hypothesis was that no difference exists among groups. False discovery rate (FDR) was controlled by adjusting p value using Benjamini-Hochberg algorithm. For DEG set, hierarchical clustering analysis was performed using complete linkage and Euclidean distance as a measure of similarity. Gene-enrichment and functional annotation analysis and pathway analysis for significant gene list were performed based on gProfiler (https://biit.cs.ut.ee/gprofiler/orth).

### Code availability

Code related to this work will be made availiable at https://github.com/baelab/PE-SB.

