## Supplementary informations for "AI-generated small binder improves prime editing"

This file includes:

Supplementary Tables

Supplementary Figures

#### Supplementary Tables

#### Supplementary Table 1. List of MC small binders

| Rank | Length | RMSD | pLDDT | i-PAE | Distance | RNF2_FC | HEK3_FC | Protein sequence |
|---|---|---|---|---|---|---|---|---|
| 1 | 150 | 1.451 | 0.919 | 6.567 | 45.438 | 0.35 | 2.42 | SKEELIKQVETAHKEAEAAFLAKFAAAKNLADFQAAYDDYLTSYASFVYSLVSSEKDEELRQAIINKAFLLLLSSTDKLFDRIEKEYGPEWLKAYLERLKELAEKYKPLVSEEDWETAEKFLAEGGEKQKRGVMLLSVYAFMSKAKEASE |
| 2 | 130 | 1.342 | 0.949 | 6.6 | 9.9801 | 0.97 | 0.89 | KEEEIKKLDEIWDQMLLGLALYEYGVEVVETASEDVETQEEALETALSLVEIASELIKPAAEETGVTKLLDILKETAEKLKEALAKLKADPSKLPEIRKELKKFLEEARKKVKEAYEEVSAVLEKKKKEL |
| 3 | 50 | 1.381 | 0.896 | 6.695 | 9.8813 | 1.24 | 1.25 | EEEIIERKAREAGRLIAERLKEAVPEASEEDLEEAGRLAAEWLRELLLAE |
| 4 | 51 | 0.983 | 0.896 | 6.876 | 10.306 | 2.86 | 4.03 | MSEREKKLEEYTEEIAKKVQEEKGISKEDAEKLKELVKEWIKKELAEIEAK |
| 5 | 98 | 1.273 | 0.913 | 6.919 | 51.458 | 1.46 | 1.92 | SMEEEIEKRKEELAEERGREVRERYLTELGLPEEEAERAGEQARRATEDLYEKVLEAFKKVEGEINEEILEDIEETTRLAFDHFLEERFREERERREA |
| 6 | 148 | 1.302 | 0.905 | 6.995 | 10.089 | 1.43 | 1.64 | KWEEVKELIKEAEKHLKKALDILEELAKDPSNIEEAYKKIVEELEKAMKALDKALELVETLTKEQQEEFAKLLGEMLAEVLKRFRRLLRMGPQAHKALLEIAKLIKEENSPVKTFMEALLRVFLLVLKALGAPEEEIKTVEEALEILS |
| 7 | 125 | 1.049 | 0.889 | 6.998 | 9.8248 | 1.59 | 1.92 | MEKLEELKKEFIEEAKEIAKVMDKYARELEEKYGLDREKAERAYKELKDRFLEIMKEISEELAELFKEAAEEIDKLDISPLGKYLALIALVEILIQKLKELGEEEKAEKLKELKEKVLKEILEEA |
| 8 | 60 | 1.187 | 0.899 | 7.147 | 10.026 | 1.13 | 1.33 | MEELLKLLEEVVEKAKKELVEKQKKLREEGKLEELKEFSENAWEEFQKLVLEGLSEAFAE |
| 9 | 111 | 1.415 | 0.899 | 7.159 | 10.418 | 0.63 | 1.19 | MSTEKRIEEDEKKVEDFFEKYYPFFKKKAEALGKEYKAAEAAGDKEKLKAIEEAIKKEEKELEEKLKDRKDAMVIIALYRIANAARESEKDLEDAMKSFLEWIKEQIRQDY |
| 10 | 89 | 1.337 | 0.916 | 7.177 | 10.167 | 1.21 | 0.78 | ETQKIAEEIKDGILKGLKGAHEALKLLLEDPSKREEALALLDEEIAKAKNKTAALLAGIWYMDEVIKENPGNKIAVELKDYLQSLLEES |
| 11 | 61 | 0.744 | 0.888 | 7.188 | 10.327 | 0.94 | 1.11 | EEEELERDNEILWRILRESDAANLTVAAERIAEELGWDLAAVTAALTSLQQQIDEKIEAEE |
| 12 | 105 | 1.326 | 0.891 | 7.247 | 47.486 | 0.62 | 1.22 | GRIEKINEVKEKLNEAFEEVLKALKEDPSLTEEEALKLLFEKIKERLGEISEGAAKLIKEIIEKVDEVLKKNEPELSIEERLKTIKEIIELMLDALKKQFKEEEA |
| 13 | 147 | 0.739 | 0.929 | 7.261 | 9.9054 | 0.90 | 1.36 | SSLEELKKKLKELREKLAPYYYKADKYLEEAIKLLKEKAKEADEEEAKKLKELIAKLEELRKELDRVIWLDLHLSALSKTGEAFLKALAKLKEELKKLKEEAKKLGLEEEAKLLEKAIEALKELEKKFKELKPLIEEIHELVDKIEE |
| 14 | 66 | 1.151 | 0.888 | 7.283 | 10.272 | 1.98 | 1.61 | NNGEENRKFAQEFQEGLKEAAAKSKTKSPEEVKELLVKKFKEEGNEDLQRAAELFLSHIIKLMKEE |
| 15 | 66 | 1.325 | 0.888 | 7.337 | 10.28 | 1.73 | 1.50 | DLEVTPEMLAKARARVAETLQARIAEIRALSPEERAKKAAELEANWEEIIGKEFLRAMTEVLEEAE |
| 16 | 75 | 1.249 | 0.89 | 7.349 | 9.8981 | 0.44 | 1.06 | VEELRARLEELQAWIAERVTTVLKELLATPEGRELAKEILTAEAEKWAEEKYGPEWGEMARAIVEAQWDLIEESE |
| 17 | 95 | 1.199 | 0.902 | 7.354 | 10.01 | 1.03 | 1.61 | AEEELERRRREIERVLSRLKATVALAYGLLTTEEGKKFFEENYEDIIKYAEETLKEAEAVLEDPVVTPEEREELEREIKRLKELLEKLKKEREEK |
| 18 | 77 | 0.821 | 0.884 | 7.433 | 9.8853 | 0.70 | 0.81 | MAKEAAKNYANLVFAQMEALGDLSGASEEQIKKLKEAAKKVGEELKKKMVSAGIPEKEAEEFAKAFVETFTKLLEAK |
| 19 | 135 | 1.334 | 0.885 | 7.463 | 10.444 | 2.65 | 2.89 | SIKEEVEKLFNELFKKIEENEEAKKQYEEAKETLKEFIEKVKEEGGSEELAKEAGEKIVEHFEEISKILSDIHQKIWDLIESLSPEDRQKAIRLLVNLILDKFKEYAVTAISMLKDEKVRELLIGKLFLAVLTLI |
| 20 | 78 | 1.332 | 0.868 | 7.496 | 10.152 | 1.63 | 1.61 | MEKVKAMKEALKELMEEAIKIREEKKDDEENREKIQQELLDKIITTTTELIASLDPSAEKGAKEQLTKYFEEILSKLR |

### Supplementary Table 2. List of MCA small binders

| Rank | Length | RMSD | pLDDT | i-PAE | Distance | RNF2_FC | HEK3_FC | Protein sequence |
|---|---|---|---|---|---|---|---|---|
| 1 | 87 | 1.074 | 0.916 | 5.757 | 45.2101 | 0.97 | 3.64 | MEERMERLRKRRKEKRKEEEKKEEEENKYFNEKATELSETLLKEGLEETEKKLEEIAKEEKMTPEMLKDLKTILESILEWTKKEIEK |
| 2 | 105 | 1.303 | 0.891 | 6.465 | 30.0549 | 1.56 | 2.83 | MEERMRRLAERMREKRKEEEELEKLKTELLSEMTKEALKEFSEKYAKGEMKIEELPELVKKIAKELAAKLKDVKDKEKIEKLFEELVPEAEKTVR MIVKHVLSQL |
| 3 | 95 | 1.23 | 0.896 | 6.553 | 15.8114 | 1.52 | 3.67 | AEEEERERERERRRKLAEGLKRLGKRAKERREKEEKEELEETRENAPKLRAEALERLAKEKGAEIAAASAAGASDEELEKLIAQGMLEVVREIFEE |
| 4 | 116 | 0.824 | 0.894 | 6.621 | 44.8209 | 1.76 | 1.89 | SALDAELAAAKAAAAAVEKELAAWTKELAKKISEKVLEELKKNKEEVQKRLEEARKEGLKKGKVEELEQEELFKIVGELVKKLVKELEPELKAQ EAKKTAAAAKVRAVYLKKFAAK |
| 5 | 133 | 1.313 | 0.899 | 6.639 | 10.1237 | 0.76 | 1.22 | MEEDEKFWRILSQKIVEVLQAYEEYKENGDKEKLEKALKEAEKTIKELTKPENREAAIEAFKTLIDTAKEIIELSEKREELEEK ARQLLRAGKEEEAAKYKEKAKELEKEIREKAKKMQEKLKEIAEEQEKKR |
| 6 | 106 | 0.984 | 0.885 | 6.696 | 49.3932 | 0.90 | 1.94 | MKEEMKKWLEEAKKTPLYKSLKRLGKRIKARREEEEKKEQEYTKELAEEVKKRLKEKAPELREKVQELERKMREEGVPEEEREQALEDLIN QLILEISKEVVKEKE |
| 7 | 130 | 0.988 | 0.89 | 6.782 | 13.3843 | 1.41 | 2.28 | MEEVLERFKKRLIEELSKVSDELEKEAYPKILAAAAAARKAQTPLHAELKKAKAKLAEYKAKGDKKKAEEAAKK MRAVLGQIAKATRAAREAGEKKLNEELNKKIKEKVEELTKELAEEWKKEQLKKLLE |
| 8 | 147 | 1.315 | 0.91 | 6.805 | 14.8915 | 0.95 | 1.28 | SREELKARFERYAARRRAERPLREREEEADALIDTLRLGVDRAAAALAALPAAKALSPELEREVRKEFYIAVNLAE QAAALAAKFAAELTAEGDTALAARLAAAVRAAAARMAGLLAEALAREREEGDEVAAARLAAALARAQAILG |
| 9 | 104 | 0.823 | 0.878 | 6.814 | 21.1757 | 1.57 | 2.14 | MEEEERREREKRWERLRENVKKAREELEKKEEKEEKEFNEKIKKAMEKALEEAKKKPLEERKNYEKLLTEVLSKTFSEVFKDKKLAEETTEVVIKAILEMLKSQ |
| 10 | 115 | 1.45 | 0.896 | 6.837 | 59.5849 | 1.40 | 3.72 | MSDPELKERMKRLSERLRKLRKEREKEEEKEKKAFIEELNAALALYRAVLEERRPEIRALLAAGELEAARAAQRAAREEALAAALSILTREENK LTLEMLAEYIEKLMAEEEAAR |
| 11 | 83 | 1.123 | 0.872 | 6.853 | 20.8132 | 1.59 | 3.81 | MKEELKKKLLEWYLSQLPEEDREKLTPEQIKEYSEKLFNMIWSKIEEIEKEEKKKEEEEEKKRREERRERLKELGKKVKEKRE |
| 12 | 117 | 1.396 | 0.892 | 6.942 | 17.2844 | 2.46 | 4.53 | MEPPEEIKEGMERLRKRLRKKREEEEKKELEYLKELAKITKEAVERFVKVEAKGRKEIRKAEEEGNKEKIEEAKKKREEEIKKIAEEAVKKDKEFSLED WMALFNGLLSLIRQQMES |
| 13 | 88 | 1.286 | 0.853 | 6.996 | 9.7954 | 0.89 | 1.03 | EEEERRKRLAERMRREGLIRETKEEIEDLEENAPLLAEKLSEAIISQIKLTEEQKEKLKKLAKERVEKAIEEYKKKLEEKTKKELEEE |
| 14 | 138 | 1.039 | 0.898 | 7.011 | 10.0432 | 0.51 | 0.89 | MSMEEEEKKREEELKERLWKEAEEFKKEAKEILKKYLEFKKSLSKEKYEKAKKQAKKWEEILELLEKAAEALKTEGL EAAKEYLKKAFALLGENIGKKVYSAVKDIMENEHPEDGIALAFAFLRAAAEIIYKKEKEEE |
| 15 | 147 | 1.128 | 0.88 | 7.04 | 47.2134 | 0.76 | 1.47 | RVEKLAKALKEELEKMVEKIKENLKKKAEKYKKEIEELEKKQPEIDKLLEEAKKLKEEGKDKEKAEKYLREVLAVRA RIARLKAEAKEAEEEIKQIDKNLKEKKEEIDKKIEEAAKRLYELSLKGDEKAFVEELNKTFSELLSLIFE |
| 16 | 124 | 1.432 | 0.879 | 7.048 | 12.4033 | 1.13 | 2.36 | SSAARAAERARIEAIVARRREERKKKEEEEKKKEEEKRKEAEKLKKEVDEEIAKIIEETKKRVEEALKEGREKALEVAREGAREMILKR LEYLSKLVGEEEVEENLPAILKLVDEYLEKLLAEE |
| 17 | 102 | 0.916 | 0.872 | 7.061 | 10.313 | 2.38 | 2.69 | KFEERLKEFLEELVEETSKLLEELDAEVNAAVEPLRKARTQALNKAFKAEMEGKEEEAKKLKEEAKALEEKAKKLEAEKKKEAKKKLEEKIKEILEKKLEES |
| 18 | 96 | 1.359 | 0.877 | 7.089 | 10.1872 | 1.05 | 1.03 | SEKEELWEKFKKRVVERMSEKLDPLFDELEKIESEIEALKNRAIRYKKLDKEKAEEYRKKAEEAEKKLEEKKKELAEKAKEIVEEIAKEMAEELKK |
| 19 | 140 | 1.122 | 0.89 | 7.112 | 10.086 | 2.98 | 5.56 | MEEEKKREEERKKEEEERIEKGMKRIAERMRARREEEEKKERERAKELKEEVEKLFEEFKKHKEEASKKVNEIIEER LKALKAAKAAGDKEAAEKAKEESYKKLIEFVKELSESFRESKLKEEIEAIIKGIFDLLVREEA |
| 20 | 94 | 1.385 | 0.85 | 7.272 | 23.8243 | 2.37 | 3.06 | MEEGWKRLRENLRKKREEEEKKEQEETKKLGEELLKFLKELLEKLSKLSIEEKKEYVKEEAEKLSKDIEKKTDLEKGKLLFLKYVLKLIEENIG |
| 21 | 107 | 1.363 | 0.852 | 7.283 | 10.4203 | 3.19 | 3.92 | SLEEKLKEKKKELLKERREYEEKAEKLESIASLTEEVLEKAAEVILEYYEEDIAKVTGMPREEAREKIREIMEKKYEELKEKREKA AKEAAEYKAKAKEVLRKIRKM |
| 22 | 122 | 0.866 | 0.871 | 7.288 | 35.5342 | 1.46 | 3.03 | MKEREELEKRMKRLRERIKERQKVEEEVEKELAEARKEAVEEFKKKAEAMKEEIKKAVEEAEDEEEAREYRKEKLAKLIADTLL KIEEKEKELYKEKLGIEIELFEEDIKEAARRAAELLIN |
| 23 | 82 | 1.47 | 0.853 | 7.318 | 9.8934 | 2.81 | 4.44 | MSEEEEKRKIEEGMRRLRENLKKKRKEEEEKEKKETEELAKLAKEIVEEAKKELKAEGKWKNKEEIWEAFKKKILEKFEKLE |

#### Supplementary Table 3. List of FMCA small binders

| Rank | Length | RMSD | pLDDT | i-PAE | Distance | RNF2_FC | HEK3_FC | Protein sequence |
|---|---|---|---|---|---|---|---|---|
| 1 | 114 | 1.052 | 0.898 | 6.249 | 9.0442 | 0.67 | 1.72 | LEELLEKIKEQFEKLKKILEKKVKEKMDEAVAKQKAAAEKEKEEKVAKLEKELKELKEKLKEGKVTEKE VNMKRLALNSVKKEYENLKKELEETKEETVKETLKLILEYIKSKL |
| 2 | 136 | 0.72 | 0.911 | 6.36 | 8.9027 | 0.83 | 2.08 | SEEEERRKRREEGMERLRKRLKEKRKELEKEEEEEEKRRREALLEAVELLLEAGREAENVREGAELW AERLLERFKELAAREDVEEALELTLEQIRLAELLLLWYAEERGDEETAEHIRRVARELRERAKALAAAA |
| 3 | 85 | 0.854 | 0.901 | 6.5 | 8.4479 | 1.06 | 1.19 | SAEIKEALERIVQNRKEREAKKKEEEKKKVKENAKKTLAEAKAKYPKYYPLLKAAIDKAEKAGQDP ELAAKTMAKQLEEFEKKVR |
| 4 | 123 | 1.285 | 0.893 | 6.504 | 9.0348 | 0.94 | 1.86 | SDRTPGMLRLAERRRALEAYRAKIEEETTKKVKEEIERERKEFLERLKKAKNLEEKEEIIEEMEEEAAKLL AESEKLRAEAAKGNLNTPENAELLVKAEALDTLGNARLALAEVARKFLEEVR |
| 5 | 122 | 1.296 | 0.887 | 6.517 | 9.0026 | 0.51 | 1.47 | SSMERLRENMKAREAAQAAADAIEAKATGAALRAALARLAATDPAGLLAAAREGPEALRAFIRARAGA DLAAAAAAVAAAMVALGAIPASAAAAVAADLEATLLEGLVLVFQWIVQQLEEAE |
| 6 | 134 | 1.082 | 0.912 | 6.582 | 8.7307 | 0.21 | 0.78 | SHMKERLERLRKNRIIQRTKIEAQEIVDGLKTAYEVLADYLSKKLEEKGYEEIAEEVRKKAEKLAEEHSKA LTAEAEAKIKEIEKQKTVEEAEKLAEAAIAELKKAAKEKMKAAREELLKLAKEGVEKLKEEKR |
| 7 | 104 | 1.36 | 0.88 | 6.597 | 8.6352 | 1.95 | 2.14 | GWERLRERRKKREEEEKKREEETQKFLKEAMKKALEKLKESEEEAEKYREEIEKEAEKIKKEAEKAKDN PEKKAELDDKADDYLAAATTISLVLSLAKEEVEKE |
| 8 | 70 | 1.354 | 0.872 | 6.829 | 9.0255 | 0.33 | 0.69 | MERYKKREEEEEKEEKAENEKQAKLAAKALKEATAKVKKIHEELGLNQEEIWKLFQEEFLKKMQEMLAKE |
| 9 | 101 | 0.936 | 0.913 | 6.879 | 8.9094 | 0.46 | 0.50 | DLEKLREELEKELEELEKEEPEIEKAMEELRRLIREKGELDDELLEKILKYAERLAEALQKAIKLEKEDSP EYKEIREKLEKKLKKVGLVIFEYYLEKKKL |
| 10 | 79 | 0.992 | 0.865 | 6.934 | 8.9223 | 0.19 | 0.44 | EFEEIWERFIERLREELAKLEEELWPEIEAAETAALAGNREEREKIEKELLEKLKPKYLEVVKKVKEEIAKEWEERRKK |
| 11 | 134 | 1.308 | 0.897 | 6.971 | 9.7184 | 0.08 | 1.14 | RRELIEKARERVEKMYAKMIQQAVDDFLASLAEERARNPAVRERIQEAMRLLKEDPEKGRALLTELVT ERFTKTAAAQRKKMDAEVKAEIAAEKDKVYAEAMAKEWERLNADFPTDVDNLVRLMVDWLVQQLLE |
| 12 | 101 | 0.976 | 0.886 | 6.973 | 9.12 | 1.29 | 1.31 | DDEKTLFQAMGYLIRYAYGESEENRDKALEYFLKYMPKDPERAKKLFEELAKKLTEIFKEVKKEEEELK AKLLAEGVRDAEEIASLEGAKKFAEKLKELGL |
| 13 | 96 | 1.423 | 0.889 | 7.003 | 8.8734 | 0.81 | 1.31 | KETAERRRKEREKEEEEETKRRKEEAEKAVKLVIEESKPYIKEMLAALKAGDLEKAKAYLEKLKAKIEE VGAKMTEFAANIMKLVNDWIVKELLSE |
| 14 | 91 | 0.894 | 0.874 | 7.005 | 9.0327 | 0.56 | 1.17 | MERFEKIEKEEQKETEIMNKLVKELVELTLERMAALPKTSVEEQKELLEEVMEEAREEVIAKHKEELDE DMIETLKSLVELFSEYVKKLLD |
| 15 | 115 | 1.298 | 0.891 | 7.053 | 9.2986 | 0.86 | 1.44 | HMLEEEKKWKEEALKLLGEIAAYLRLIYSEEEREKMWLEIAKLIADFLYASELPFGSEEVKKVMEELKK KLKEYKEKAAEIGPVPEEAIERAEKAVRELIEKAKELMAKSEEAEE |
| 16 | 131 | 0.858 | 0.888 | 7.075 | 9.691 | 0.60 | 1.14 | MEEIKKRMERLSKRLKKKREEKEKALKEFNKKINELIKKNMEKLLKELKPLGKKAKEAKTEEERKKIL EEIKEKIKKAEEELKKEIEKIAEEFKKKYPEFAEEFETLEESYKELAKYLKELYIEFIEKVFN |
| 17 | 111 | 1.258 | 0.898 | 7.075 | 8.9894 | 0.51 | 1.06 | MEEREKKEREELRKEGEKLKEVFKDVITLAGAVQKQSSPSDTPSDVFRRTKAELAAIAADPAAAAA AREAAAADPEVLAILEKFLADPEKAETAITLLDLLVKWFELVESQ |

#### Supplementary Table 4. Primer sequences for qPCR

| Gene | Direction | Sequence |
|---|---|---|
| 18s rRNA | Forward primer | GTAACCCGTTGAACCCCATT |
| | Reverse primer | CCATCCAATCGGTAGTAGCG |
| MLH1 | Forward primer | ATGCCCACCAGATGGTTCGT |
| | Reverse primer | CCCTTTGTTGTATCCCCCTCC |

#### Supplementary Table 5. Primer sequence for targeted deep sequencing

| Gene | Direction | Sequence | |
|---|---|---|---|
| HEK3 | Forward primer | ACACTCTTTCCCTACACGACGCTCTTCCGATCT | aaacgcccatgcaattagtc |
| | Reverse primer | GTGACTGGAGTTCAGACGTGTGCTCTTCCGATCT | ccagccaaacttgtcaacc |
| HEK4 | Forward primer | ACACTCTTTCCCTACACGACGCTCTTCCGATCT | ctcccttcaagatggctgac |
| | Reverse primer | GTGACTGGAGTTCAGACGTGTGCTCTTCCGATCT | aacggagacacacacacagg |
| RNF2 | Forward primer | ACACTCTTTCCCTACACGACGCTCTTCCGATCT | ccagcaatgtctcaggctgt |
| | Reverse primer | GTGACTGGAGTTCAGACGTGTGCTCTTCCGATCT | GCCAACATACAGAAGTCAGGAA |

| Gene | Direction | Sequence | |
|---|---|---|---|
| HEK3_OT1 | Forward primer | ACACTCTTTCCCTACACGACGCTCTTCCGATCT | TCCCCTGTTGACCTGGAGAA |
| | Reverse primer | GTGACTGGAGTTCAGACGTGTGCTCTTCCGATCT | CACTGTACTTGCCCTGACCA |
| HEK3_OT2 | Forward primer | ACACTCTTTCCCTACACGACGCTCTTCCGATCT | TTGGTGTTGACAGGGAGCAA |
| | Reverse primer | GTGACTGGAGTTCAGACGTGTGCTCTTCCGATCT | CTGAGATGTGGGCAGAAGGG |
| HEK3_OT3 | Forward primer | ACACTCTTTCCCTACACGACGCTCTTCCGATCT | TGAGAGGGAACAGAAGGGCT |
| | Reverse primer | GTGACTGGAGTTCAGACGTGTGCTCTTCCGATCT | GTCCAAAGGCCCAAGAACCT |
| HEK4_OT1 | Forward primer | ACACTCTTTCCCTACACGACGCTCTTCCGATCT | GGCATGGCTTCTGAGACTCA |

| | | | |
|---|---|---|---|
| | Reverse primer | GTGACTGGAGTTCAGACGTGTGCTCTTCCGATCT | CCCTTGCACTCCCTGTCTTT |
| HEK4_OT2 | Forward primer | ACACTCTTTCCCTACACGACGCTCTTCCGATCT | TTTGGCAATGGAGGCATTGG |
| | Reverse primer | GTGACTGGAGTTCAGACGTGTGCTCTTCCGATCT | GAAGAGGCTGCCCATGAGAG |
| HEK4_OT3 | Forward primer | ACACTCTTTCCCTACACGACGCTCTTCCGATCT | TTTCCACCAGAACTCAGCCC |
| | Reverse primer | GTGACTGGAGTTCAGACGTGTGCTCTTCCGATCT | CCTCGGTTCCTCCACAACAC |
| RNF2_OT1 (Cas OFFinder) | Forward primer | ACACTCTTTCCCTACACGACGCTCTTCCGATCT | TACTGTGTAAGCGGAATGGG |
| | Reverse primer | GTGACTGGAGTTCAGACGTGTGCTCTTCCGATCT | CTGACGTTTCTGAAGGTTGC |
| RNF2_OT2 (Cas OFFinder) | Forward primer | ACACTCTTTCCCTACACGACGCTCTTCCGATCT | GTTTACCATGATACCTGTGCCCT |
| | Reverse primer | GTGACTGGAGTTCAGACGTGTGCTCTTCCGATCT | TGGCTGTGAGGATGTGTAGG |
| RNF2_OT3 (Cas OFFinder) | Forward primer | ACACTCTTTCCCTACACGACGCTCTTCCGATCT | CACCAGGTATCTGTCTTGACAATC |
| | Reverse primer | GTGACTGGAGTTCAGACGTGTGCTCTTCCGATCT | TCAGCATTCTATGACCTTGATGTC |
| RNF2_OT4 (Cas OFFinder) | Forward primer | ACACTCTTTCCCTACACGACGCTCTTCCGATCT | GTGAGCAGCCCTACAGAGAG |
| | Reverse primer | GTGACTGGAGTTCAGACGTGTGCTCTTCCGATCT | ATTTGGGCAGGGACACACATC |
| RNF2_OT5 (Cas OFFinder) | Forward primer | ACACTCTTTCCCTACACGACGCTCTTCCGATCT | GGGCTGGGAGGGAAAGTTAT |
| | Reverse primer | GTGACTGGAGTTCAGACGTGTGCTCTTCCGATCT | GCCCAGAAAGCTTCCTTACC |

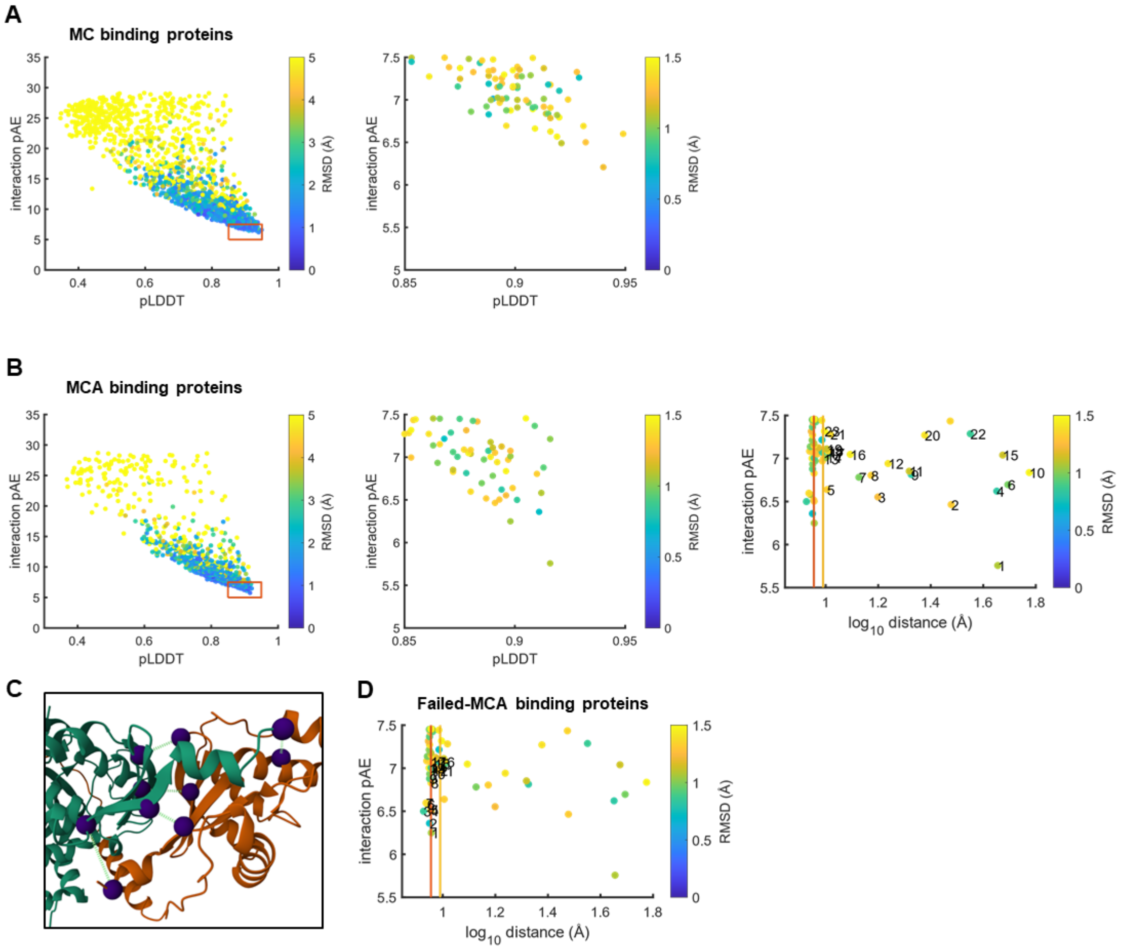

**Figure S1.**

(A) Distribution of parameters for MC binder proteins (left). Total number was 1978. Filtered distribution using the threshold values (middle), 7.5 for interaction pAE, 0.85 for pLDDT, 1.5 Å for RMSD. Filtered MC binder proteins by the threshold value, 9.7724 Å (yellow line), are marked with the 20 top rank numbers (right). Red line is at the original distance, 9.0157 Å, same as in Figure 1D. (B) Distribution of parameters for MCA binder proteins (left). Total number was 821. Filtering process is same as in (A). Selected MCA binders are marked with the 23 top rank numbers (right). (C) Selected five atom pairs (purple) to calculate the average atom-pair distance of the Cα atoms of the residues (i.e. interface distance), L540, L743, Q542, C756, N739 in MLH1-C and I688, I853, N683, E705, Q861 in PMS2-C. (D) Filtered failed-MCA binder proteins by the threshold. Failed-MCA binder proteins are marked with the 17 top rank numbers.

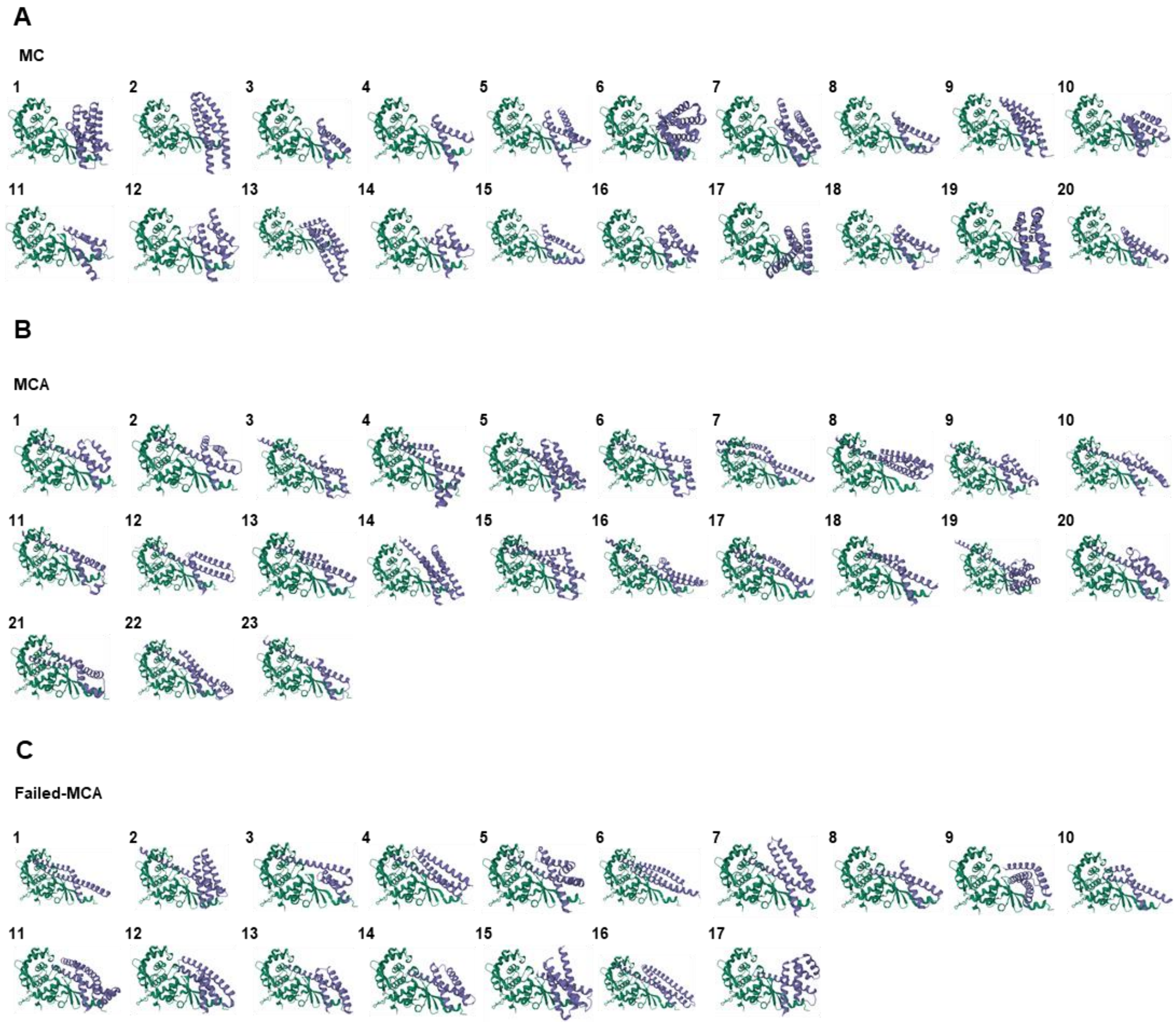

**Figure S2.**

Dimeric structures of C-terminal domain of MLH1 (MLH1-C) and MLH1 small binders, predicted by AlphaFold 2 integrated in the RFdiffusion program, for MC binder proteins (A), MCA binder proteins (B) and failed-MCA binder proteins (C). MLH1-C is colored in green, and MLH1 small binders are colored in blue.

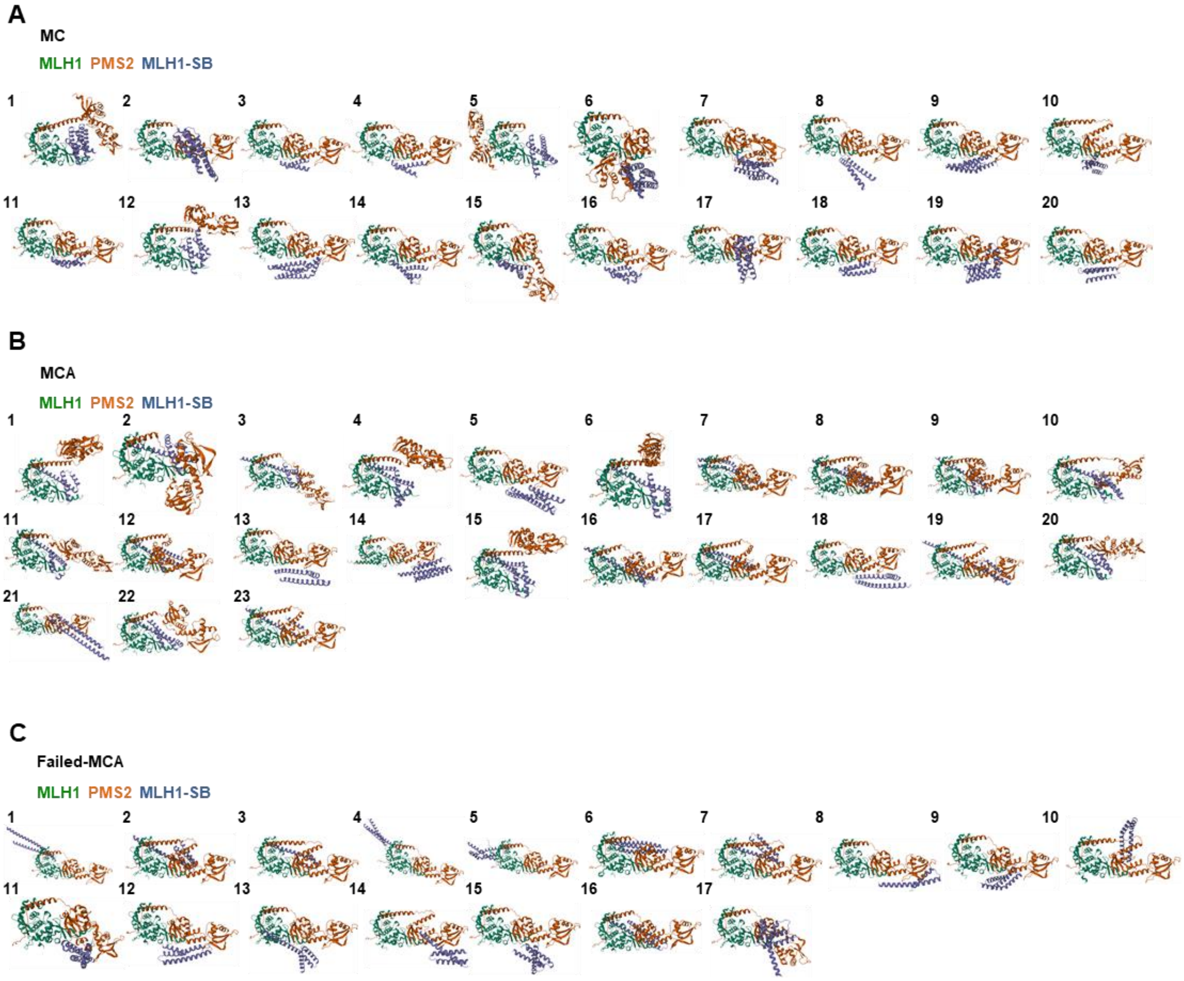

**Figure S3.**

Trimeric structures of C-terminal domain of MLH1 (MLH1-C), C-terminal domain of PMS2 (PMS2-C) and small binders, predicted by AlphaFold 3, for MC binder proteins (A), MCA binder proteins (B) and failed-MCA binder proteins (C). MLH1-C is colored in green, PMS2-C is colored in orange, and MLH1 small binders are colored in blue.

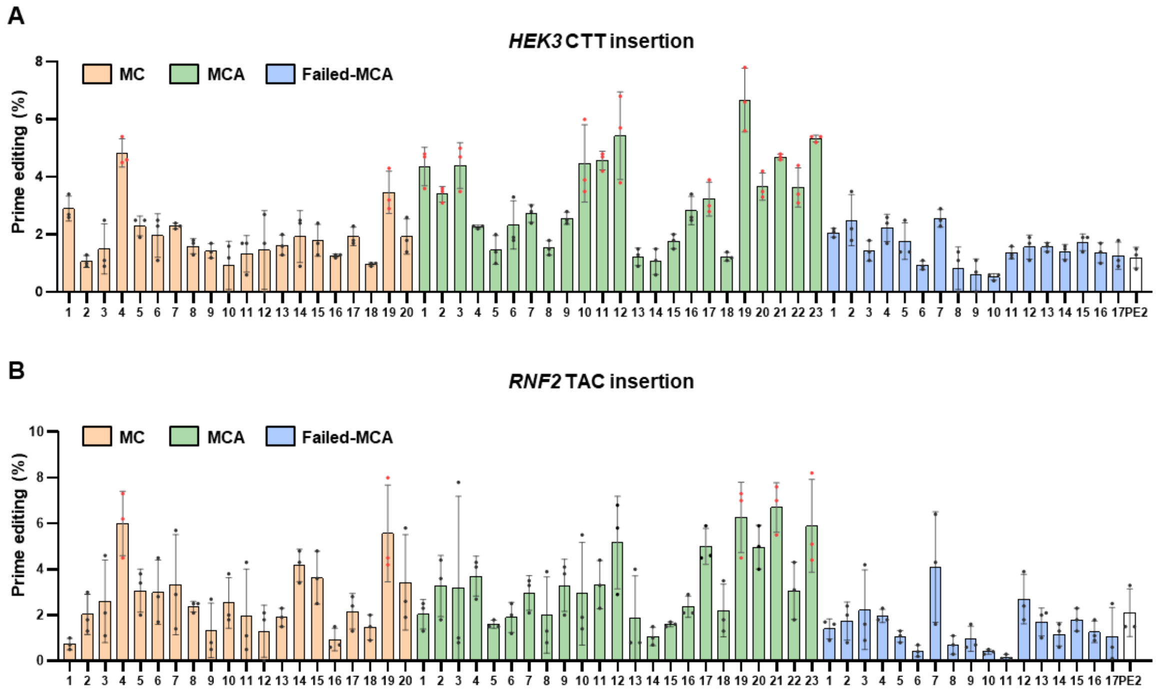

**Figure S4.**

(A-B) Prime editing efficiency for *HEK3* CTT insertion (A) and *RNF2* TAC insertion (B) using designated MLH1 small binders. MLH1 small binders that demonstrate more than a 2.5-fold increase in prime editing efficiency are marked with red dots.

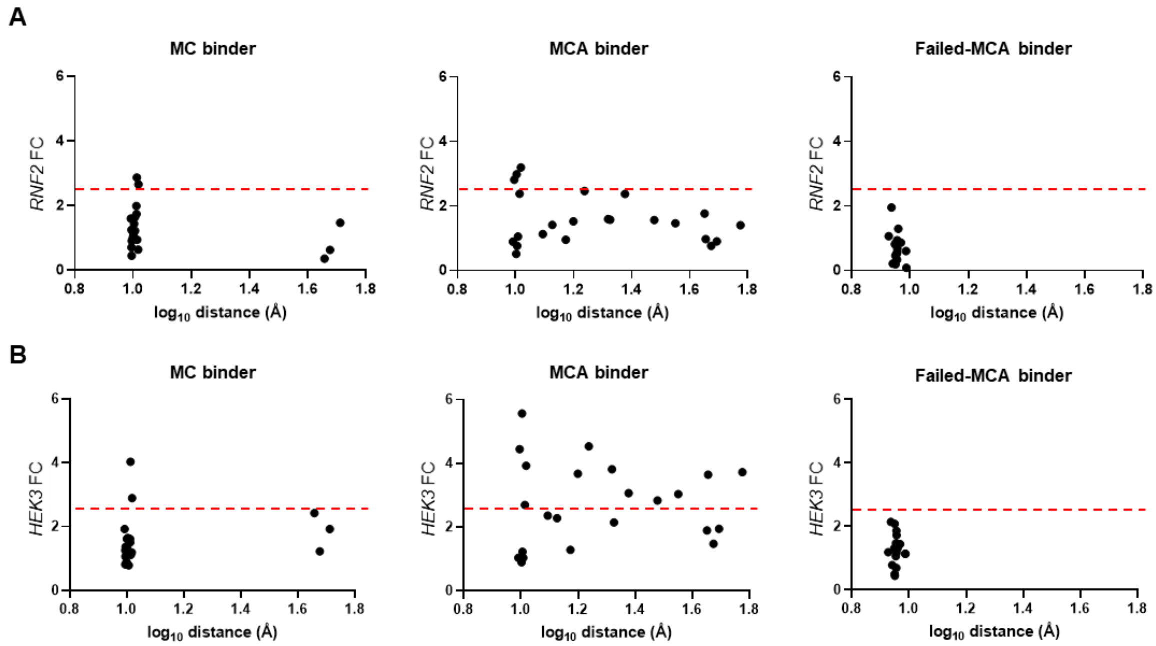

**Figure S5.**

Fold change distribution of prime editing efficiency for *RNF2* TAC insertion (A) and *HEK3* CTT insertion (B) using MC binders (left), MCA binders (middle) and failed-MCA binders (right). A 2.5-fold increase is highlighted with a red dotted line.

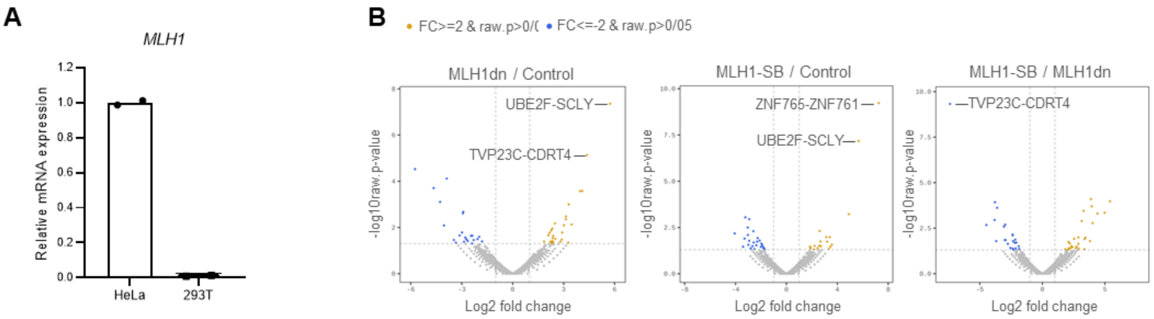

C **MLH1dn / Control**

| source | term_id | term_name | adj.p.value |
|---|---|---|---|
| GO:BP | GO:0050789 | regulation of biological process | 0.19068 |
| GO:BP | GO:0050794 | regulation of cellular process | 0.19068 |
| GO:BP | GO:0065007 | biological regulation | 0.19068 |
| GO:BP | GO:0051716 | cellular response to stimulus | 0.19623 |
| GO:BP | GO:0007154 | cell communication | 0.19873 |
| GO:BP | GO:0023052 | signaling | 0.19873 |
| GO:BP | GO:0007188 | adenylate cyclase-modulating G protein-coupled receptor signaling pathway | 0.20536 |
| GO:BP | GO:0007165 | signal transduction | 0.20536 |
| GO:CC | GO:0016020 | membrane | 0.22642 |
| GO:BP | GO:0050896 | response to stimulus | 0.24364 |

D

**MLH1-SB / Control**

| source | term_id | term_name | adj.p.value |
|---|---|---|---|
| GO:MF | GO:0004305 | ethanolamine kinase activity | 0.09354 |
| GO:MF | GO:0004103 | choline kinase activity | 0.09354 |
| GO:BP | GO:0006657 | CDP-choline pathway | 0.14449 |
| GO:BP | GO:0006646 | phosphatidylethanolamine biosynthetic process | 0.14449 |
| GO:BP | GO:0046337 | phosphatidylethanolamine metabolic process | 0.18797 |
| GO:BP | GO:0006656 | phosphatidylcholine biosynthetic process | 0.18797 |
| GO:MF | GO:0043168 | anion binding | 0.19257 |
| GO:MF | GO:0016773 | phosphotransferase activity, alcohol group as acceptor | 0.19257 |
| GO:BP | GO:0006629 | lipid metabolic process | 0.20110 |
| GO:MF | GO:0016301 | kinase activity | 0.21539 |

E

**MLH1-SB / MLH1dn**

| source | term_id | term_name | adj.p.value |
|---|---|---|---|
| GO:BP | GO:0008593 | regulation of Notch signaling pathway | 0.08294 |
| GO:BP | GO:0007219 | Notch signaling pathway | 0.13164 |
| GO:BP | GO:0016310 | phosphorylation | 0.13496 |
| GO:MF | GO:0097367 | carbohydrate derivative binding | 0.13989 |
| GO:BP | GO:0000082 | G1/S transition of mitotic cell cycle | 0.14821 |
| GO:MF | GO:0004693 | cyclin-dependent protein serine/threonine kinase activity | 0.15024 |
| GO:MF | GO:0032559 | adenyl ribonucleotide binding | 0.15024 |
| GO:MF | GO:0097472 | cyclin-dependent protein kinase activity | 0.15024 |
| GO:MF | GO:0030332 | cyclin binding | 0.16221 |
| GO:BP | GO:0044843 | cell cycle G1/S phase transition | 0.16366 |

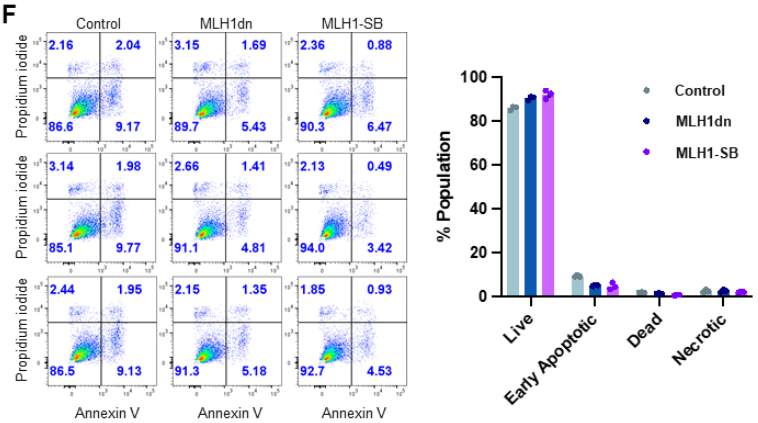

**Figure S6.**

(A) Relative mRNA expression levels of *MLH1* in HeLa and HEK293T cells, Bars represents mean values for n=2 independent biological replicates. (B) Volcano plots depicting the differential gene expression from MLH1dn over control (left), MLH1-SB over control (middle), and MLH1-SB over MLH1dn (right). The x-axis represents the log2 fold change, and the y-axis represents the -log10 of the raw p-value. Genes with significant upregulation (Log2FC $\geq$ 2, raw $p < 0.05$) are highlighted in orange, and significantly downregulated (Log2FC $\leq$ -2, raw $p < 0.05$) genes are highlighted in blue. (C-E) Gene Ontology (GO) enrichment analysis of differentially expressed genes for MLH1dn over Control (C), MLH1-SB over control (D), and MLH1-SB over MLH1dn (E). Adjusted p-values for each term are provided to indicate significance. (F) Flow cytometry result of AnnexinV-propidium iodide (PI) staining assay of HeLa cells transfected with puromycin resistant gene expression vector (control), dominant negative MLH1 expression vector (MLH1dn), and MLH1 small binder expression vector (MLH1-SB) (left) and bar graph of each part of populations (right). The bottom left quadrant (Annexin V−/PI−) indicates live cells, the top left quadrant (Annexin V−/PI+) indicates necrotic cells, the bottom right quadrant (Annexin V+/PI−) shows early apoptotic cells, and the top right quadrant (Annexin V+/PI+) shows late apoptotic or dead cells. Numbers in each quadrant reflect the proportion of cells within these states. Bars represent mean values, and error bars represent the S.D. of n = 3 independent biological replicates.

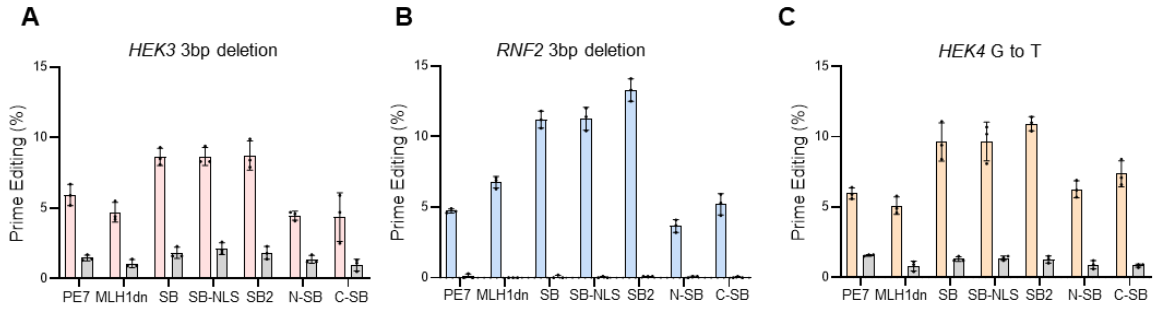

**Figure S7**

(A-C) Prime editing efficiency and unwanted mutations at designated loci and mutation by PE7, PE7 with MLH1dn (MLH1dn), PE7 with MLH1-SB (SB), PE7 with MLH1-SB-NLS (SB-NLS), PE7 with both MLH1-SB and MLH1-SB-NLS (SB2), PE7 linked to MLH1-SB at N-terminal (N-SB) or C-terminal (C-SB) of PE7 at *HEK3* (A), *RNF2* (B), and *HEK4* (C) in HeLa cells. Bars represent mean values, and error bars represent the S.D. of n = 3 independent biological replicates.

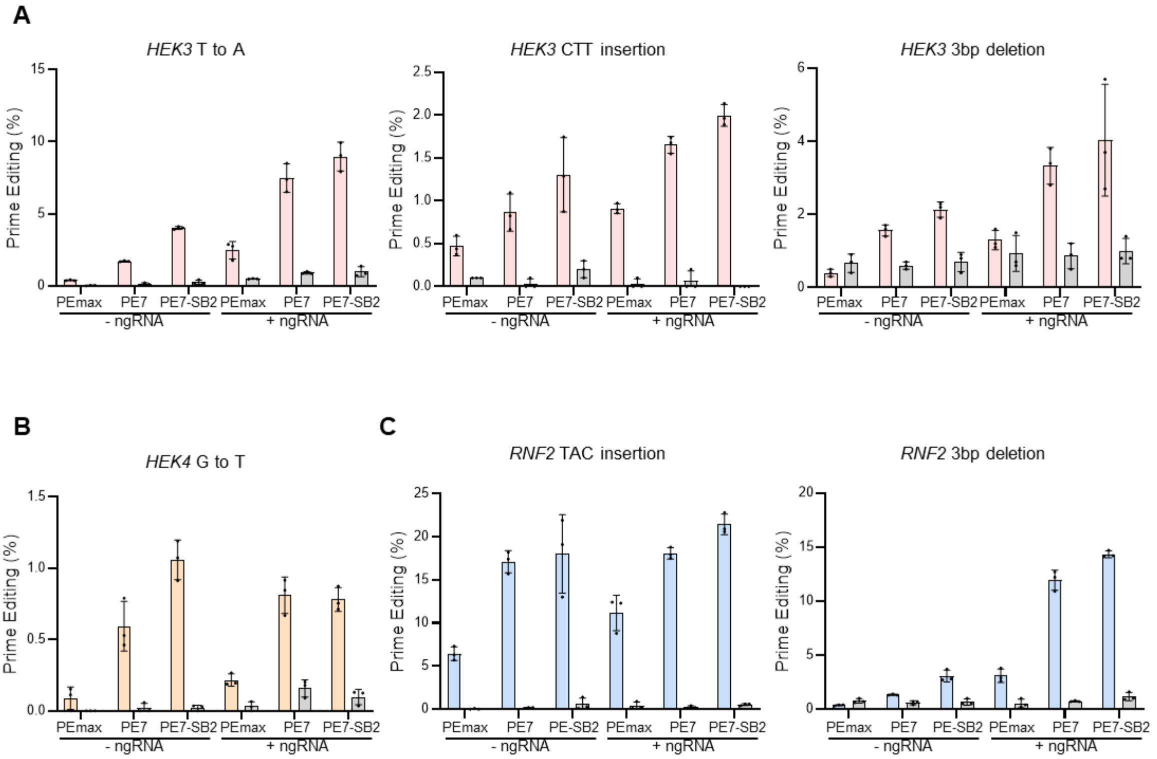

#### Figure S8

(A-C) Prime editing efficiency and unwanted mutations for designated mutation by PEmax, PE7, and PE7-SB2 with or without nicking gRNA (ngRNA) at *HEK3* (A), *HEK4* (B), and *RNF2* (C) in hiPSCs. Bars represent mean values, and error bars represent the S.D. of n = 3 independent biological replicates.

**A**

| Target | Spacer |
|---|---|
| HEK3 | GGCCCAGACTGAGCACGTGA |
| OT1 | CACCCAGACTGAGCACGTGC |
| OT2 | GACACAGACCGGGCACGTGA |
| OT3 | AGCTCAGACTGAGCAAGTGA |

| Target | Spacer |
|---|---|
| HEK4 | GGCACTGCGGCTGGAGGTGG |
| OT1 | TGCACTGCGGCCGGAGGAGG |
| OT2 | GGCTCTGCGGCTGGAGGGGG |
| OT3 | GGCATCACGGCTGGAGGTGG |

| Target | Spacer |
|---|---|
| RNF2 | GTCATCTTAGTCATTACCTG |
| OT1 | GGTATCTAAGTCATTACCTG |
| OT2 | GTCATCCTAGTCATTTACTG |
| OT3 | GTAATATTAGTCATTACCGG |
| OT4 | GTCATCTGAGGCATTAACTG |
| OT5 | GTAATCTGAGTCATTTCCTG |

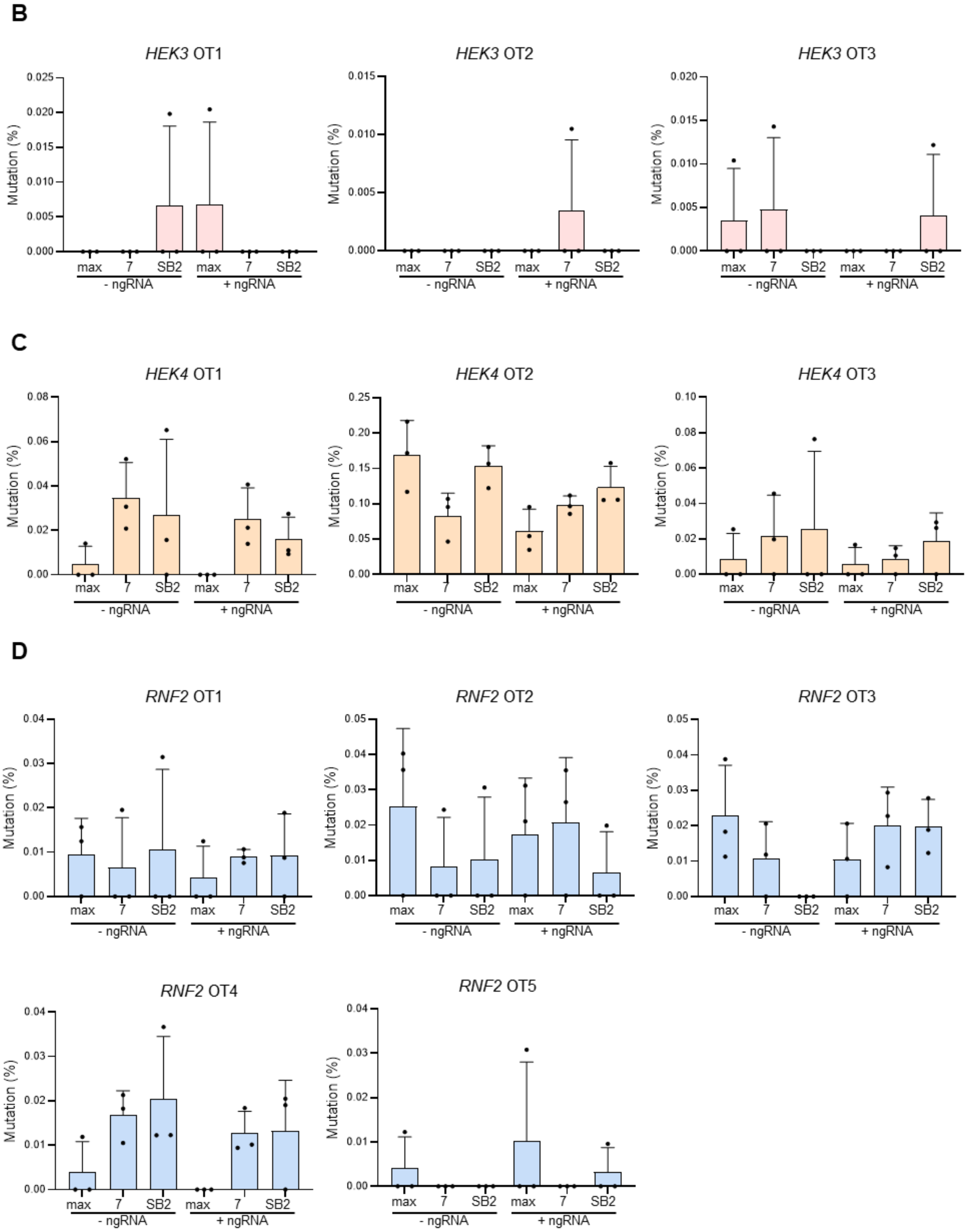

**Figure S9**

(A) Sequence information of off-targets of *HEK3* (left), *HEK4* (middle) and *RNF2* (right).

Unmatched bases of the off-target spacers are colored in red. (B-D) Off-target mutations of *HEK3* CTT insertion (B), *HEK4* G to A substitution (C), and *RNF2* TAC insertion (D) by PEmax (max), PE7 (7), and PE-SB2 (SB2) with or without nicking gRNA (ngRNA) in HeLa cells. Bars represent mean values, and error bars represent the S.D. of n = 3 independent biological replicates.
